# Global analysis of expression, maturation and subcellular localization of mouse liver transcriptome identifies novel sex-biased and TCPOBOP-responsive long non-coding RNAs

**DOI:** 10.1101/2021.01.14.426753

**Authors:** Christine N. Goldfarb, David J. Waxman

**Author notes:** Correspondence: Dr. David J. Waxman, Dept. of Biology, Boston University, 5 Cummington Mall, Boston, MA 02215, **Email**.

## Abstract

While nuclear transcription and RNA processing and localization are well established for protein coding genes (PCGs), these processes are poorly understood for lncRNAs. Here, we characterize global patterns of transcript expression, maturation and localization for mouse liver RNA, including more than 15,000 lncRNAs. PolyA-selected liver RNA was isolated and sequenced from four subcellular fractions (chromatin, nucleoplasm, total nucleus, and cytoplasm), and from the chromatin-bound fraction without polyA selection. Transcript processing, determined from normalized intronic to exonic sequence read density ratios, progressively increased for PCG transcripts in going from the chromatin-bound fraction to the nucleoplasm and then on to the cytoplasm. Transcript maturation was similar for lncRNAs in the chromatin fraction, but was significantly lower in the nucleoplasm and cytoplasm. LncRNAs were 11-fold more likely to be significantly enriched in the nucleus than cytoplasm, and 100-fold more likely to be significantly chromatin-bound than nucleoplasmic. Sequencing chromatin-bound RNA greatly increased the sensitivity for detecting lowly expressed lncRNAs and enabled us to discover and localize hundreds of novel regulated liver lncRNAs, including lncRNAs showing sex-biased expression or responsiveness to a xenobiotic agonist ligand of constitutive androstane receptor (Nr1i3). Integration of our findings with prior studies and lncRNA annotations identified candidate regulatory lncRNAs for a variety of hepatic functions based on gene co-localization within topologically associating domains or transcription divergent or antisense to PCGs associated with pathways linked to hepatic physiology and diseases.

## Background

Since the discovery of more than a thousand novel, poly-adenylated long non-coding RNAs (lncRNAs) in mouse and human cells [1], lncRNAs have increasingly been shown to play key roles in gene regulation and disease states, including liver disease [2-4]. LncRNAs typically have 5’ caps, are transcribed by RNA polymerase II, and have polyA tails, and the DNA from which they are transcribed can have promoter-like or enhancer-specific histone modifications [1, 5]. Many lncRNA genes display striking patterns of developmental, condition-dependent and tissue-specific expression, which enables them to serve as condition-specific regulators of diverse biological processes [6]. LncRNAs can regulate cellular functions at multiple levels, including epigenetic modification and chromatin remolding, transcriptional regulation, alternative splicing and mRNA translation [7-9]. For example, the lncRNA Xist, which is crucial for X-chromosome dosage compensation in female cells in eutherian mammals, introduces a repressed chromatin state marked by extensive histone-H3 K27me3 across one of the X-chromosomes, leading to X-inactivation and Barr body formation [10, 11], while the oncogenic lncRNA HOTAIR promotes cancer metastasis in part by silencing HOXA genes by promoting K27-trimethylation and K4-demethylation of histone-H3 [12]. In the liver, lncRNAs have been linked to liver fibrosis through their effects on glucose metabolism (LincIRS2) [13] and hepatic stellate cell regulation (H19, Meg3, HOTTIP) [14-16]. However, the biological functions and mechanisms of action of the vast majority of lncRNAs expressed in liver and other tissues are unknown.

Many lncRNAs are preferentially localized in the nucleus, where they can be visualized by single molecule RNA fluorescence in situ hybridization (smFISH) [17]. A few dozen lncRNAs have thus been characterized and show diverse patterns of expression, ranging from one or two distinct nuclear foci per cell to many individual RNA molecules throughout the nucleus and/or cytoplasm [18]. Multiplex error-robust FISH enables a higher throughput visualization of lncRNA and mRNA transcripts, though probes still need to be designed individually for each RNA of interest [19, 20]. LncRNAs can also be localized by RNA-seq analysis of subcellular RNA fractions. In prior studies from this laboratory, poly-adenylated RNA was isolated from nuclei purified from fresh liver tissue and sequenced to identify liver-expressed lncRNAs [21, 22]. Other studies using rRNA-depleted RNA from human hepatocellular carcinoma cell lines also showed nuclear enrichment of lncRNAs but not RNAs coding for protein-coding genes (PCGs) [23]. While some cytoplasmic lncRNAs have been described [24], a majority of well-studied lncRNAs appear to function primarily in the nucleus.

Within the nucleus, lncRNAs may be nucleoplasmic, may be associated with the nuclear matrix or other structures, or may be tightly bound to chromatin, where they can interact directly with chromatin modifying complexes and regulate transcription. LncRNAs that bind to specific chromatin modifying complexes have been identified by RNA immunoprecipitation, although there are concerns about promiscuity and non-specific binding [25]. Related technologies have enabled the discovery of the specific RNAs, proteins and genomic regions that interact with individual lncRNAs [26], but this approach cannot readily be implemented on a global scale to study the thousands of lncRNAs expressed in a given cell line or tissue. However, by fractionating nuclei using a high urea buffer containing salts and detergent, RNAs that are tightly bound to chromatin can be separated from RNAs that are soluble in the nucleoplasm, enabling the characterization of several thousand lncRNAs enriched in the insoluble chromatin fraction from human cell lines and mouse macrophages [27, 28].

We previously identified 15,558 lncRNAs expressed in mouse liver under a variety of biological conditions [29], a subset of which were characterized with regard to their tissues-specific expression patterns, epigenetic states, and regulatory elements, including nearby regions of chromatin accessibility and liver-specific transcription factor binding [22]. These liver-expressed lncRNAs include lncRNA genes that are responsive to xenobiotic exposure [21, 30] or to pituitary growth hormone secretory patterns [22], a key factor regulating sex-biased gene expression in the liver [31, 32]. Co-expression network analysis in Diversity Outbred mice identified sex-biased lncRNAs whose expression is inversely correlated with that of oppositely sex-biased PCGs, suggesting they have important negative regulatory actions in mouse liver [29]. We also found that many lncRNAs induced or repressed in xenobiotic-exposed rat liver are closely linked to xenobiotic dysregulation of pathways involving fatty acid metabolism, cell division and immune responses [33]. However, key information regarding subcellular localization is lacking for the vast majority of these liver-expressed lncRNAs, which complicates efforts to determine whether they have regulatory functions in the cytoplasm, nucleoplasm or when bound to chromatin.

Here, we use RNA-seq to characterize the expression patterns for a set of 15,558 liver-expressed lncRNAs and compare them to more than 20,000 PCGs across four subcellular fractions and under four different biological conditions. We identify genes whose transcripts are present at significantly different levels between the cytoplasm and the nucleus, and for nuclear transcripts, between the nucleoplasm and a chromatin-bound fraction. We find an unexpectedly strong enrichment of thousands of liver-expressed lncRNAs in the chromatin fraction, including lncRNAs that respond to endogenous hormonal factors or external chemical exposure, many expressed at too low a level for discovery by traditional RNA-seq analysis of whole liver tissue or even in purified liver nuclei. Our analysis of these rich datasets gives new insights into the maturation of hepatic lncRNA transcripts, and integration of our findings with prior work enabled us to identify lncRNAs that serve as strong candidates for future investigations of lncRNA function in liver biology and disease.

## Results

### RNA-seq analysis of liver subcellular fractions

We sought to identify liver-expressed genes whose transcripts are differentially enriched between the cytoplasmic and nuclear compartments. Frozen liver tissue, from untreated male and female mice, and from mice exposed to TCPOBOP, an CAR agonist ligand that induces or represses several hundred genes [21, 34], was homogenized under conditions expected to preserve nuclear membrane integrity. RNA was isolated from washed nuclei and from the cytoplasmic lysate, and nuclei were used to further separate and then purify soluble, nucleoplasmic RNA and RNA tightly bound to chromatin after extraction with high salt buffer and urea **(Fig. S1)**. RNA purified from each subcellular fraction (cytoplasm, nucleus, nucleoplasm, chromatin-bound fraction) was analyzed by qPCR to determine the localization and regulated expression of select sex-biased or TCPOBOP-responsive marker genes. Elovl3 showed strong, male-biased expression in the cytoplasmic, nuclear and nucleoplasmic fractions of untreated liver (**Fig. 1A**). TCPOBOP induced Elovl3 in the chromatin-bound fraction in male liver, and in all four fractions in female liver, which largely abolished its sex-dependent expression. The primary transcript, pre-Elovl3 RNA, showed highest expression in the chromatin-bound fraction. pre-Elovl3 RNA was induced ≥10-fold in all three nuclear-derived fractions by TCPOBOP treatment in both male and female liver, indicating that TCPOBOP stimulates Elovl3 gene transcription (**Fig. 1B**). The differential enrichment of mature Elovl3 vs. pre-Elovl3 RNA in each subcellular fraction validates the separation of the fractions. Further validation was obtained by examining Cyp2b10, which showed female-biased expression in untreated liver and was strongly induced by TCPOBOP (up to 300-fold) in both sexes (**Fig. 1C**). The lncRNA Neat1 (lnc14746) was exclusively found in the nuclear and chromatin-bound fractions (**Fig. 1D**). Furthermore, Xist (lnc15394), which is only expressed in female cells, showed similar expression levels in the nuclear, nucleoplasmic and chromatin-bound fractions and was absent from cytoplasm (**Fig. 1E**), as expected [18].

**Fig. 1.**
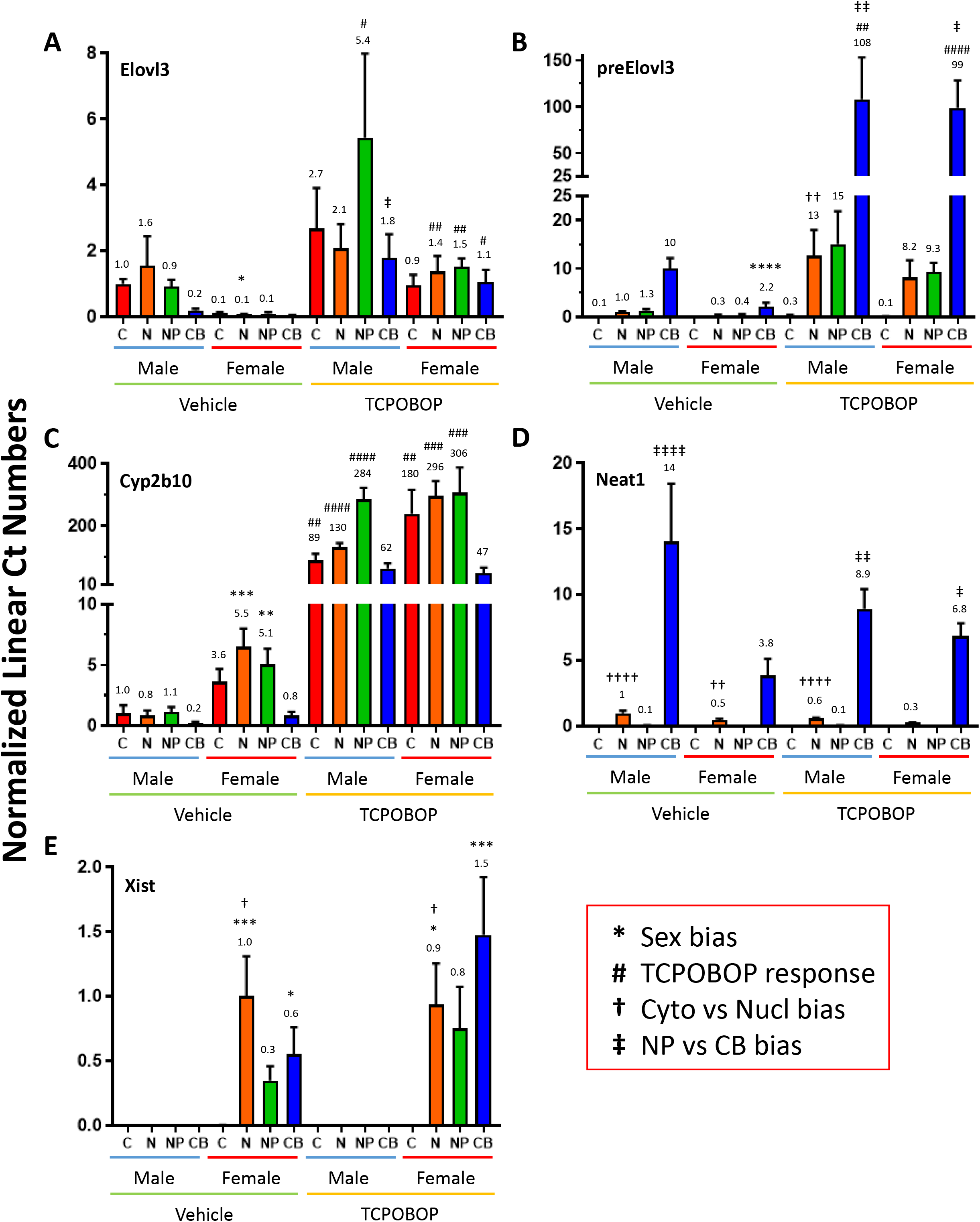
qPCR analysis of liver subcellular fractions using select marker genes. Expression of each gene was determined by qPCR across the cytoplasm (C), nucleus (N), nucleoplasm (NP) and chromatin-bound (CB) fractions. Data shown are relative expression levels (values above each bar) as mean values ± SEM for n=4 mice per biological condition: vehicle treated male and female mice, and TCPOBOP-treated male and female mice. (**A**) <u>Elovl3</u>: male-biased expression seen in vehicle-treated mice is largely lost following TCPOBOP treatment, which induces Elovl3 in both male and female liver. (**B**) <u>PreElovl3</u> was assayed using qPCR primers that span an intron/exon boundary to amplify unspliced transcripts, which were significantly enriched in the chromatin-bound fraction after TCPOBOP exposure in both sexes. (**C**) <u>Cyp2b10</u> analysis validated the strong TCPOBOP induction response, and also female-biased expression in the basal state. (**D**) The highly chromatin-bound <u>lnc14746/Neat1</u> (significant in 3 of the 4 biological conditions) validated the separation of the nucleoplasmic and chromatin-bound fractions. (**E**) The female-specific <u>lnc15394/Xist</u> showed strong expression in all three nuclear-derived fractions. Significance was determined by one-way ANOVA with Bonferroni correction, for four separate analyses, which are specified using four different symbols (red box), as follows: p < 0.05, one symbol; p < 0.01, two symbols; p < 0.001, three symbols; and p < 0.0001, four symbols. qPCR primers are shown in Table S1A.

Next, to obtain a global view of the localization and regulation of liver RNAs, including liver-expressed lncRNA genes, we prepared RNA-seq libraries from polyA-selected RNA from each of the four fractions (n=3-4 livers per biological condition). We also sequenced the chromatin-bound fraction without polyA-selection to obtain expression data for both poly-adenylated and non-poly-adenylated RNAs, including transcripts that did not yet undergo polyadenylation. In all, we sequenced 65 RNA-seq samples representing the 5 cellular fractions under 4 different biological conditions (male and female liver, with and without TCPOBOP exposure) (**Table S1A**). These datasets were then analyzed to address questions related to lncRNA maturation and localization and regulation, as described below.

### Transcript maturity in different subcellular fractions

We used the following approach to assess transcript maturity for each liver-expressed, multi-exonic lncRNA and PCG (**Table S1D**). Reads mapping to exon collapsed (EC) regions, and separately, reads mapping to intronic only (IO) regions, were counted for each gene, and then normalized by the % exonic and % intronic length of the gene, respectively. The resultant normalized exonic and intronic read densities were used to calculate an intronic to exonic read density ratio, IO/EC (**Table S1F**. For a transcript that is completely unspliced (i.e., a primary, immature transcript), RNA sequence reads will be spread equally across the entire gene length, and the IO/EC ratio will equal 1; and for a transcript that is fully spliced, the intronic read count, and hence the IO/EC ratio, will equal 0. Thus, lower IO/EC ratios are associated with increased transcript processing (increased RNA maturity).

We found that IO/EC ratios progressively decreased in going from the chromatin-bound to the nucleoplasmic and nuclear fractions, and then on to the cytoplasm, with the decrease in ratio being much greater for PCG than for lncRNA gene transcripts (**Fig. 2A**; adjusted p-value < 0.0001 for Cyto vs NP, and for NP vs CB, for both lncRNAs and PCGs). Thus, transcripts in the chromatin bound fraction are the least spliced/most immature, and as RNAs transition from the chromatin bound state through the nucleoplasm and then on the cytoplasm, there is a progressive increase the extent of RNA maturation. Median IO/EC ratios were 4-11-fold lower for PCGs than for lncRNAs in the cytoplasmic, nuclear and nucleoplasmic fractions (for PCGs and lncRNAs, respectively, median cytoplasmic ratio = 0.0032 and 0.036; median nuclear ratio = 0.027 and 0.12; and median nucleoplasmic ratio = 0.015 and 0.089; all significant at adjusted p-value < 0.0001). Thus, lncRNA splicing is less efficient/less complete than PCG splicing, consistent with reports that at least some incompletely spliced lncRNAs are biologically active (see Discussion). Median IO/EC ratios were much higher, and were not significantly different between PCGs and lncRNAs in the chromatin-bound fractions (IO/EC = 0.20 and 0.23 for polyA-selected PCGs and lncRNAs, respectively; IO/EC = 0.31 and 0.34 for non-polyA-selected PCGs and lncRNAs, respectively). Thus, the chromatin-bound fraction contains many more unspliced or partially spliced transcripts, in particular in the non-polyA selected fraction (**Fig. 2B, Fig. 2C, Fig. S2**).

**Fig. 2.**
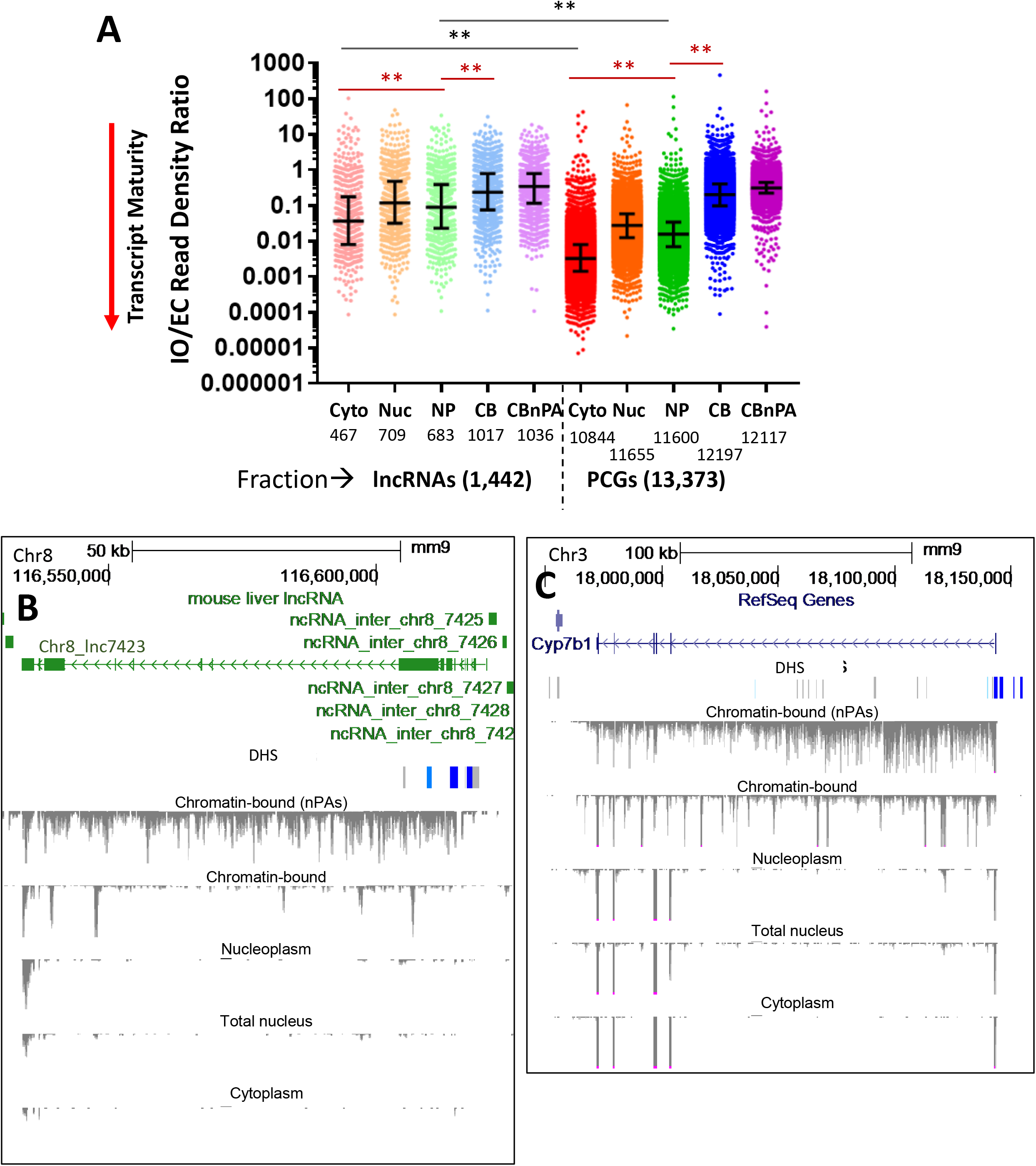
Transcript maturity across subcellular fractions determined by IO/EC read density ratio analysis. (**A**) Distributions of IO/EC read density ratios for individual genes in vehicle-treated male liver, calculated from the weighted normalized read density values for the intronic only (IO) and the exonic collapsed (EC) reads for each of 1,442 multi-exonic lncRNAs (left) and 13,737 multi-exonic PCGs (right). IO/EC ratios displayed are mean values for n=3 livers. The number of genes expressed in each subcellular fraction (see Methods) is listed below each column: cytoplasm (Cyto), nucleus (Nuc), nucleoplasm (NP), chromatin-bound (CB), chromatin bound non-PolyA selected (CBnPAs). Median values (black horizontal midline) and IQR (error bars) are marked. Black horizontal lines compare the distributions of IO/EC ratios for lncRNAs vs PCGs in the cytoplasmic and nucleoplasmic fractions (other comparisons were not performed); red horizontal lines compare distributions between the indicated fractions for lncRNAs, and separately, for PCGs (** = adjusted p-value < 0.0001). The higher IO/EC ratios apparent for nuclear compared to nucleoplasmic transcripts is due to the nuclear fraction being a composite of both nucleoplasmic and chromatin-bound RNA. An excess of normalized intronic reads (IO/EC ratios > 1) is seen for a subset of genes, most notably chromatin-bound PCGs and all five lncRNA fractions. Many of these genes are lowly expressed (very low normalized EC reads), but have short, unannotated expressed features; others have intronic regions that overlap with an exon of an expressed gene, leading to an artefactually high IO read count and IO/EC ratio. The data used to generate these graphs are found Table S1F. Fig. S2 shows similar results for vehicle-treated female liver. (**B**) and (**C**) UCSC Browser screen shot showing BigWig files of minus strand sequence reads for each of the five indicated subcellular fractions for lnc7423 (gene structure shown in green) and Cyp7b1 in untreated male mouse liver. Extensive reads seen across the gene body in the chromatin bound fraction are substantially depleted after polyA-selection (top vs second reads track); however, multiple distinct peaks within intronic regions remain. BigWig Y-axis scale: 0 to −25, except for non-polyA-selected track, which is 0 to −5 (B) or 0 to −12 (C). Both genes show male-biased expression, with many fewer sequence reads in corresponding fractions from female liver (not shown). Cytoplasm, and to a lesser extent nucleoplasm, are depleted of sequence reads for lnc7423 but not for the PCG Cyp7b1, where a progressive increase in transcript maturity is apparent. These same patterns were seen in all three biological replicates. DHS, DNase hypersensitivity sites, indicating open chromatin. DHS showing significantly greater accessibility in male liver are marked in blue [100]. Also see Fig. S2.

### Differential enrichment of transcripts in subcellular fractions

We sought to identify RNAs showing differential intracellular localization, as indicated by significant differential expression between subcellular fractions. RNA-seq samples were normalized across samples based on total reads mapping to exonic features (exon collapsed read counts), and differential expression analysis was then to identify transcripts significantly enriched at high stringency in cytoplasmic vs nuclear fractions, and separately, nucleoplasmic vs chromatin-bound fractions, and in the chromatin-bound fractions with vs without polyA selection (**Tables S2A-S2C**).

Many more lncRNAs were significantly enriched in nuclear RNA as compared to cytoplasmic RNA (n=748 vs n=64; 11.7-fold difference) (**Table S2A**). Forty of the 64 cytoplasmic-biased lncRNAs were antisense lncRNAs, several of which were antisense to very highly expressed PCGs whose exons line up with a majority of lncRNA sequence reads found on the opposite strand. The cytoplasmic reads associated with these antisense lncRNAs may arise from incomplete second strand degradation during RNA-seq library preparation, and could therefore be artefactual. Seven of the 24 other cytoplasmic-biased lncRNAs were responsive to TCPOBOP treatment and two showed sex-biased expression (adjusted p-value < 0.05) (discussed further below). The high preponderance of nuclear-biased lncRNAs contrasts with a more equal distribution between cytoplasmic-biased (n=1,460) and nuclear-biased PCG transcripts (n=1,767) **(Fig. 3A; Fig. S3A, Fig. S3B)**. The median expression level was significantly lower for the subcellular fraction-biased lncRNAs compared to the correspondingly biased PCGs (**Fig. 3B**; 3.9-fold lower for cytoplasm and 51-fold lower for nucleus). Further, the subcellular fraction expression ratio ranged much higher for the nuclear-biased transcripts than for the cytoplasmic-biased transcripts, most notably for the lncRNAs **(Fig. 3C)**. The strong nuclear enrichment of many lncRNAs (median nuclear to cytoplasmic ratio = 12.5-fold (IQR, 6.5 to 22.3) suggests those transcripts are not exported to the cytoplasm, or are exported but then rapidly degraded in the cytoplasm. The much lower cytoplasmic bias seen for PCGs (median cytoplasmic to nuclear ratio = 2.1-fold (IQR, 1.89 to 2.45) may reflect their ongoing transcription to generate a robust basal level of nuclear transcripts, which would effectively dampen the cytoplasmic to nuclear ratio.

**Fig. 3.**
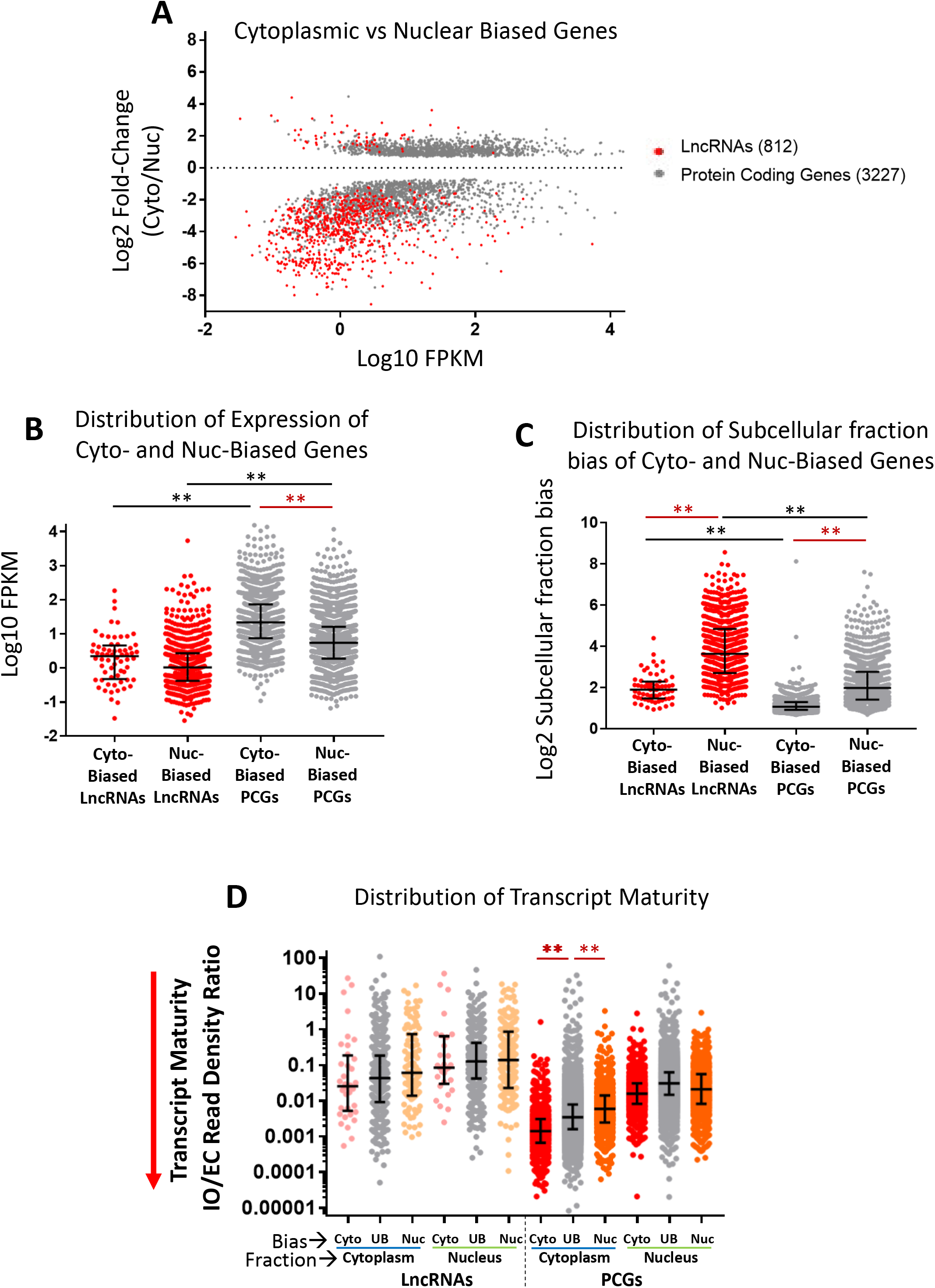
Expression, subcellular fraction bias and maturity of cytoplasmic versus nuclear transcripts. (**A**) Subcellular fraction bias (compartment bias), displayed as cytoplasmic (Cyto) to nuclear (Nuc) expression ratio, of all RNAs that show either cytoplasmic-biased (positive y-axis) or nuclear-biased transcript levels (negative y-axis) at an edgeR-adjusted p-value < 0.001 in at least one of the four biological conditions assayed. For genes showing significant compartment bias in more than one condition, data is shown for the condition with the highest FPKM value (Table S2A, columns D and E). The data points represent 1,524 cytoplasmic-biased genes (64 lncRNAs, 1,460 PCGs) and 2,515 nuclear-biased genes (748 lncRNAs, 1,767 PCGs). Data for lncRNAs and PCGs are regraphed separately in Fig. S3A and Fig. S3B. (**B**) Distributions of FPKM values, and (**C**) distribution of compartment bias values (i.e., differential expression between subcellular fractions) for the four indicated sets of subcellular fraction-enriched RNAs. The median fraction bias was 1.8-1.9-fold higher (adjusted p-value < 0.0001) for the nuclear-biased transcripts than for the cytoplasmic-biased transcripts. (**D**) Distributions of transcript maturity values (normalized IO/EC read density ratios, from Table S1F) in the cytoplasmic and nuclear fractions (“Fraction”) for multi-exonic lncRNAs and multi-exonic PCGs that show a significant cytoplasmic bias (Cyto) or nuclear bias (Nuc) (“Bias”), or that do not show a significant compartment bias (UB, unbiased). For **B, C**, and **D**, median values (black horizontal midline) and IQR (error bars) are indicated; black horizontal lines compare lncRNAs to PCGs within the same fraction, and red horizontal lines compare lncRNAs, or PCGs, between groups, as marked, with ** indicating adjusted p-value < 0.0001. In D, statistical analysis was used to compare Cyto vs UB, and UB vs Nuc, for lncRNAs and PCGs with the cytoplasmic or nuclear fractions.

Transcript maturity may in part drive these differences in expression between subcellular fractions, at least for PCGs. Thus, PCG transcripts enriched in the cytoplasm are on average more mature (lower median IO/EC ratio) than the corresponding fraction-unbiased and nuclear-enriched transcripts. Importantly, the greater transcript maturity of cytoplasm-enriched PCG RNAs was observed in both the cytoplasmic fraction, and separately, in the nucleus (**Fig. 3D**), and is associated with a significantly shorter gene length, but not a lower percentage of intronic sequence (**Fig. S3C, Fig. S3D)**. Of note, in the nucleus, transcript maturity was similar, or even higher, for nuclear-biased PCGs and lncRNAs compared to non-compartment-biased PCGs and lncRNAs (**Fig. 3D**), suggesting that other factors, such as chromatin binding, examined below, drive the nuclear bias of these transcripts.

### Widespread enrichment of lncRNAs in chromatin-bound RNA

RNA-seq analysis of nucleoplasmic and chromatin-bound RNA extracted from liver nuclei identified 3,057 compartment-biased lncRNAs, of which 3,028 (99%) were significantly enriched in the chromatin fraction (**Table S2B**). Preferential expression in the chromatin fraction was also seen for 92% of 7,719 other lncRNAs that did not meet our stringent criteria (adjusted p<0.001) for differential enrichment between fractions (**Fig. S4B, Table S2B**). Moreover, 13 of the 29 lncRNAs enriched in the nucleoplasm were antisense lncRNAs, which in several cases may be artefacts of incomplete second strand degradation during library preparation, as discussed above. In contrast, PGCs were more likely to be significantly enriched in the nucleoplasm (n=3,264) than in chromatin (n=1,420) **(Fig. 4A, Fig. S4A)**, suggesting they are rapidly transcribed and then released from their chromatin-associated transcriptional complexes. Indeed, the strength of the compartment bias was much weaker for chromatin-enriched PCG transcripts than for chromatin-enriched lncRNAs (median chromatin/nucleoplasm ratio: PCGs, 3.1 (IQR, 2.5 to 4.5), lncRNAs, 18.5 (IQR, 8.4 to 60.4) (**Fig. S4C**), consistent with the proposal that many PCG transcripts but not many lncRNA transcripts are efficiently release to the nucleoplasm following transcription. Transcript maturity was significantly higher for all classes of PCGs, but not for lncRNAs, in the nucleoplasm than in the chromatin fraction, further supporting with this model (**Fig. S4E**).

**Fig. 4.**
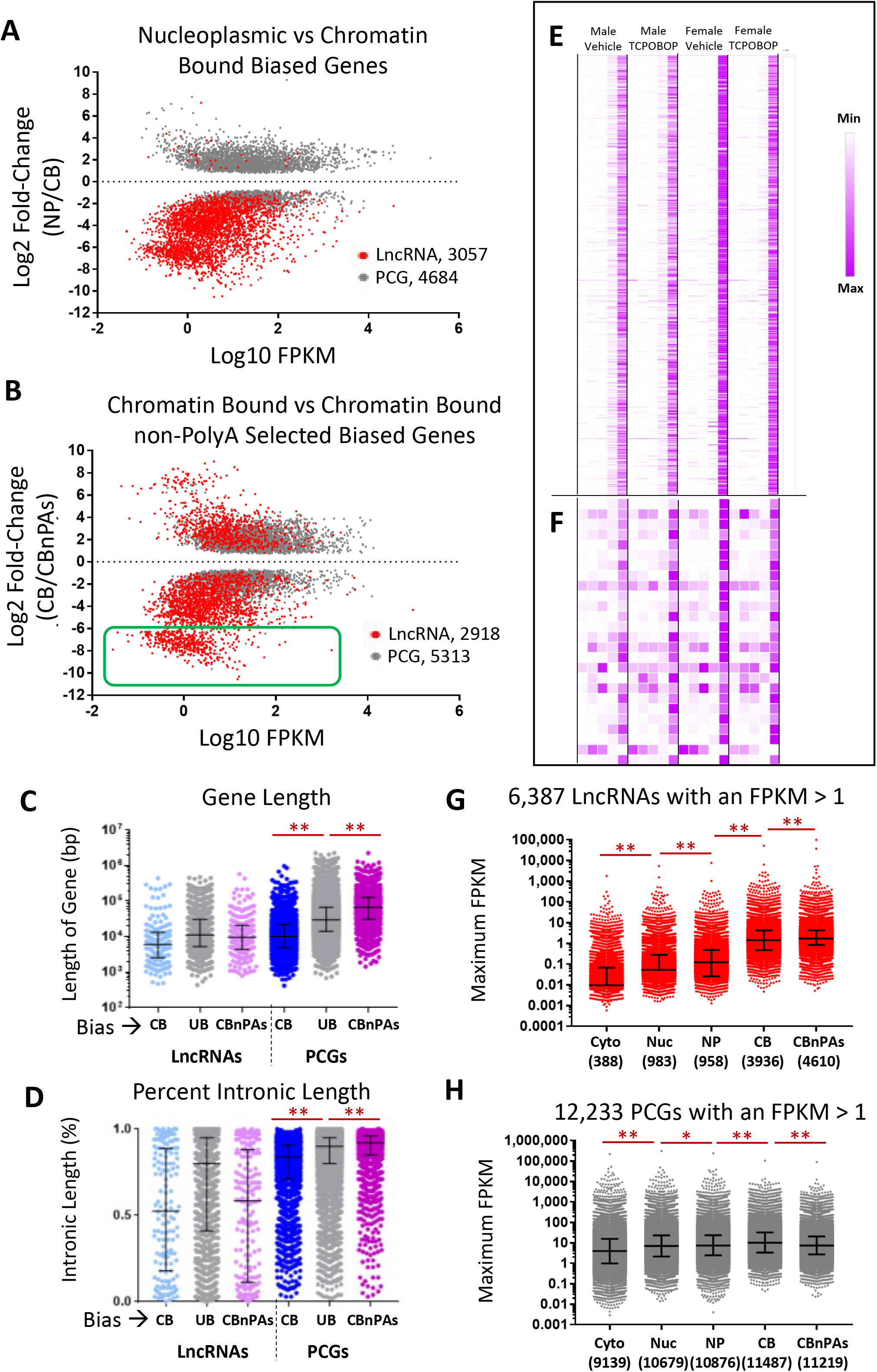
Differential expression of lncRNAs and PCGs across nuclear subcellular fractions. Subcellular fraction bias between: **(A)** nucleoplasm (NP) and the chromatin-bound (CB) fraction; or **(B)** within the chromatin-bound fraction, between polyA-selected and non-polyA selected RNA, based on an edgeR adjusted p-value < 0.001 in at least one of the four biological conditions assayed, and based on data shown in Table S2B and Table S2C, columns D and E. Gray dots are PCGs, red dots are lncRNAs. For any gene showing a significant compartment bias in more than one biological condition, data is shown for the condition with the highest FPKM value. Genes total: in **A**, 3,293 nucleoplasm-biased genes (29 lncRNAs, 3,264 PCGs) and 4,448 chromatin-biased genes (3,028 lncRNAs, 1,420 protein coding genes); and in **B**, 4,027 chromatin-biased, polyA-selected genes (844 lncRNAs, 3,183 PCGs) and 4,204 chromatin-biased, non-polyA-selected (CBnPAs) genes (2,074 lncRNAs, 2,130 PCGs). Green box, CBnPAs-biased genes with log2 fold-change < −6, which are further analyzed in **E. (C)** and **(D)**, Distributions of gene lengths (**C**) and percent intronic length (**D**) for chromatin-bound biased (CB), non-compartment-biased (UB, unbiased) and CBnPAs-biased, graphed separately for lncRNAs and PCGs; also see Table S1D, columns M-Q. Significant differences for PCGs are as marked; no significant differences were seen for lncRNAs. See Fig. S4 for corresponding data for NP-biased vs CB-biased genes, and Fig. S5 for CB-biased genes, with vs without polyA selection. (**E)** and (**F**), Normalized expression for the 506 lncRNAs (**E**) and 26 PCGs (**F**) that were very strongly CBnPAs-biased (genes from green box in **B**) across all 4 biological conditions (marked at top), for each of 5 subcellular fractions (columns from left to right: Cytoplasm, Nucleus, Nucleoplasm, Chromatin-bound, and Chromatin-bound non-PolyA-selected). See data in Table S2C, columns X-AM. Data are shown for expression of each gene (row), normalized to the highest expression of that gene in a single condition and fraction. Of note, 17 of the 506 lncRNAs show sex-biased expression (Table S3A), and 19 show TCPOBOP-responsiveness (Table S3B) in at least one subcellular fraction. (**G)** and (**H**), Distribution of expression values (FPKM) for the subsets of 6,387 lncRNAs (**G**) and 12,233 PCGs (**H**) that are expressed at FPKM > 1 in at least one of the 5 subcellular fractions. The maximum expression of the gene across the four biological conditions is graphed for each subcellular fraction. Only a subset of the lncRNAs and PCGs were expressed FPKM >1 in each fraction, as indicated by the gene count numbers below each column. Based on expression data in Tables S2A-S2C, columns X-AM. Median FPKM values (black horizontal midline) and IQR (error bars) are marked. Red horizontal lines compare lncRNAs, or PCGs, between fractions: adjusted p-value < 0.05 (*) and adjusted p-value < 0.0001 (**).

Finally, we investigated whether the nucleoplasmic and chromatin enriched PCG RNAs are enriched for different biological processes. Enriched terms for the most highly nucleoplasm-biased PCGs (subcellular fraction bias > 5 and FDR <0.05; n=914 PCGs) include transmembrane helix, secreted, extracellular matrix, cadherin, blood coagulation and immunity (**Table S2D**); while the most highly chromatin bound-biased PCGs (subcellular fraction bias > 5 and FDR < 0.05; n=755 PCGs) were most highly enriched for the terms synapse, sequence-specific DNA binding, ion channel activity, and multicellular organism development (**Table S2E**).

### Impact of polyA selection on RNA profiles

Many more lncRNAs were significantly enriched in chromatin-bound RNA without polyA selection, which includes non-poly-adenylated transcripts, than in the polyA-selected fraction (n=2,074 vs. n=844). In contrast, PCGs were more commonly enriched in the polyA-selected fraction (n=3,185 vs n=2,128) (**Fig. 4B, Table S2C, Fig. S5A-S4D)**. The greater tendency for PCGs to be enriched in the polyA-selected fraction, where they have a significantly higher transcript maturity compared to the non-polyA-selected fraction **(Fig. S5E)**, is consistent with the association of poly-adenylation with transcript maturation [35]. In contrast, the tendency for lncRNAs to be enriched in the non-polyA-selected fraction is consistent with splicing being delayed or incomplete for lncRNAs [36]. The enrichment of a subset of the chromatin-bound PCGs in the non-polyA-selected fraction was associated with a significantly longer gene length and comparatively longer introns (adjusted p-value < 0.0001, for both) as compared to chromatin-bound PCGs enriched in the polyA-selected fraction (**Fig. 4C, 5D**), consistent with these PCGs requiring longer times for completion of transcription and/or processing prior to poly-adenylation. Longer gene lengths were also seen when comparing nuclear-enriched to cytoplasm-enriched PCGs **(Fig. S3C)**, but not when comparing chromatin-enriched to nucleoplasm-enriched PCGs **(Fig. S4F)**. Finally, enriched terms for the most highly enriched genes in the polyA-selected chromatin-bound fraction (compartment bias > 5 and FDR <0.05; n=933 PCGs) include ribosomal protein, oxidative phosphorylation/mitochondria, non-alcoholic fatty liver disease, and mRNA-splicing (**Table S2F**); while the most highly enriched terms for the non-polyA-selected chromatin-bound PCGs (compartment bias > 5 and FRD < 0.05; n=776 PCGs) included nucleosome assembly [primarily histone genes, whose transcripts are not poly-adenylated [37]], metal binding/zinc finger proteins, Pleckstrin homology domain, and DNA-binding (**Table S2G**).

We observed a cluster of chromatin-bound transcripts, comprised of 506 lncRNAs and 26 PCGs, with > 64-fold higher levels in the non-polyA-selected than in the polyA-selected fraction (**Fig. 4B**, green box). All of these lncRNAs show their highest expression in the chromatin-bound, non-polyA-selected fraction across all four treatment groups (**Fig. 4E**), consistent with these being lncRNA transcripts that do not undergo poly-adenylation. Similarly, 23 of the 26 PCGs were most highly expressed in the non-polyA-selected fraction (**Fig. 4F**), including several histones RNAs, which as noted are not poly-adenylated [37]. Other PCGs in this group include the gap junction protein Gja6 and two beta-cadherin protogenes (Pcdhb11, Pcdhb21) and three zinc-finger genes (Rnf148, Zfp691, Zfp804b).

### Increased sensitivity for lncRNA detection in chromatin-bound fraction

Global analysis of lncRNA expression levels across the five subcellular fractions (**Tables S2A-S2C**) revealed a striking increase in the number of lncRNAs detected at FPKM >1 when going from the cytoplasm (n=388) to the nucleus (n=983) or nucleoplasm (n=958) to the chromatin-bound fractions (n=3,936, n=4,610). Moreover, median lncRNA expression levels increased significantly across the five fractions, with the sensitivity for lncRNA detection increasing 32-fold when analyzing chromatin-bound non-polyA RNA (median expression = 1.69 FPKM, IQR, 0.84 to 4.22) as compared to total nuclear RNA (median expression = 0.052 FPKM, IQR, 0 to 0.28) (adjusted p-value < 0.001) (**Fig. 4G**). PCGs did not show any subcellular fraction-dependent increase in expression (**Fig. 4H**).

### Discovery of sex-biased and TCPOBOP-responsive lncRNA transcripts

Given the striking enrichments of distinct sets of lncRNAs in each subcellular fraction and the increased sensitivity of lncRNA detection seen in chromatin-bound RNA, we used our datasets to discover novel regulated lncRNAs. Differential expression analysis of untreated male versus female liver identified 701 sex-biased genes, including 375 sex-biased lncRNAs and 20 other non-coding RefSeq genes (**Fig. 5A; Table S3A**). The vast majority of the lncRNAs (352/375, 94%) showed sex-biased expression in one or both chromatin-bound fractions, whereas only 68 (18%) showed sex-biased expression in the cytosol or nucleoplasm. This contrasts to a much larger fraction of sex-biased PCGs (168 of 306 genes; 55%) identified in the cytosol or nucleoplasm (**Table S3A, Fig. 5B)**. Similarly, we found that large numbers of lncRNAs were either induced or repressed by the CAR agonist ligand TCPOBOP in male or female liver (**Table S3B** and **Fig. 5C;** 1,005 lncRNAs and 131 other noncoding RefSeq genes, including 26 miRNAs). Many of these lncRNAs responded to TCPOBOP in one sex only (**Fig. 5D**, left two columns of each gene set), as was also the case for many TCPOBOP-responsive PCGs, consistent with our prior findings [21]. Furthermore, 81% of the 1,005 lncRNAs regulated by TCPOBOP were responsive in one or both chromatin-bound fractions, as compared to only 14% that responded in the cytoplasm and 22% in the nucleoplasm (**Table S3B**). Thus, large numbers of regulated lncRNAs are readily identified by RNA-seq analysis of chromatin-bound liver RNA, highlighting the advantages of using RNA-seq to analyze chromatin-bound RNA for discovery of condition-specific, transcriptionally-regulated lncRNA genes.

**Fig. 5.**
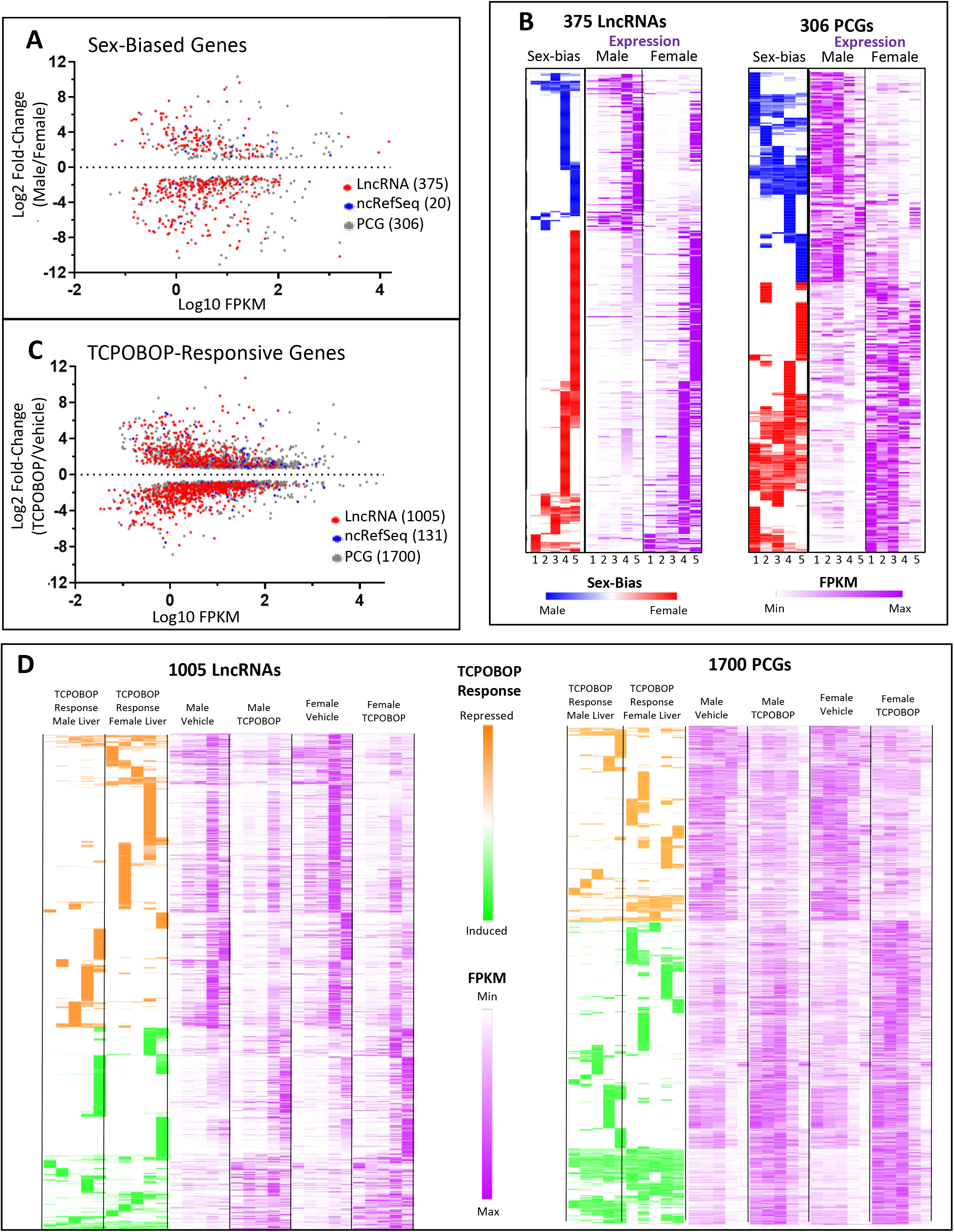
Expression and responsiveness of sex-biased and TCPOBOP-responsive genes. (**A**) Genes showing differential expression between male and female liver (edgeR adjusted p-value cutoff of 0.05) in at least one of the five subcellular fractions assayed. See data in Table S3A, columns D and E. Gene totals: 123 male-biased and 252 female-biased lncRNAs, 11 male-biased and 9 female-biased non-coding RefSeq genes, and 134 male-biased and 172 female-biased PCGs. (**B**) The sex-bias and expression in both male and female liver are shown across all 5 subcellular compartments (left to right, Cytoplasm, Nucleus, Nucleoplasm, Chromatin Bound, Chromatin Bound non-PolyA selected) (Table S3A, columns Z-AS). Expression of each gene (row) is normalized to the highest expression (or strongest sex-bias) of the gene across all fractions. (**C**) Genes showing differential expression between vehicle and TCPOBOP-treated liver (edgeR adjusted p-value cutoff of 0.05) in at least one of the five subcellular fractions assayed. See data in Table S3B, columns D and E. In **A** and **C**, for any gene that is significantly biased in more than one fraction, the fraction with the maximum FPKM and its corresponding fold-change is graphed. Gene totals: 411 up and 594 down regulated lncRNAs, 69 up and 62 down regulated non-coding RefSeq genes, and 1035 up and 665 down regulated PCGs. (**D**) TCPOBOP responsiveness and expression in both male and female liver is shown across all 5 subcellular compartments (left to right, Cytoplasm, Nucleus, Nucleoplasm, Chromatin Bound, Chromatin Bound non-PolyA selected) (Table S3B, columns AF-BS). Expression of each gene (row) is normalized to the highest expression or TCPOBOP responsiveness of that gene in a single condition and fraction. In many cases, significant differential expression was seen in only one subcellular fraction for both sex-biased genes (B) and TCPOBOP-responsive genes (D); in many cases, the same trends were apparent but lacked statistical significance due to very low expression in other fractions and/or variation between biological replicates (**Table S3A**).

### smFISH analysis of lncRNA localization

We used smFiSH (RNAScope technology; **Fig. 6A** [38]) to localize two sex-biased lncRNAs in mouse liver slices. lnc7423 **(Fig. 2B**), which shows significant male-biased expression, was visualized at several sites per cell in male liver, while in female liver, only a few cells showed expression (**Fig. 6B; Fig. S6A, Fig. S6B**), consistent with the strong, male-bias expression in several nuclear fractions seen by RNA-seq (**Table S3A**). lnc14770, a female-biased lncRNA, was detected at less than one copy per cell in male liver, but in female liver, five or more copies were seen in some cells, although many cells had only one copy (**Fig. 6C; Fig. S6C, Fig. S6D**). For both sex-biased lncRNAs, expression was almost exclusively nuclear, and appeared as focal dots, consistent with tight chromatin binding. Our RNA-seq data showed that lnc7423 is 4-6-fold enriched in the chromatin fraction in both sexes, whereas the female-biased lnc14770 only showed a significant nuclear bias only in female liver (22-fold; **Table S7**). We also visualized Cyp2b10 and its divergently transcribed (5.1 kb upstream) lnc5998, both of which are highly induced by TCPOBOP. In untreated male liver, Cyp2b10 expression was very low, with a few RNA molecules detected in the cytoplasm, while lnc5998 was essentially undetectable. Following TCPOBOP exposure, large dense clouds of Cyp2b10 RNA surrounded each nucleus, consistent with the high induction of this RNA seen by RNA-seq and its association with endoplasmic reticulum membrane-bound polysomes. Cyp2b10 showed 3-fold nucleoplasmic bias in TCPOBOP treated liver in both male and female liver (**Table S7**). Very strong induction of lnc5998 was also apparent, which in contrast to Cyp2b10 RNA, was more concentrated in nuclei, consistent with its highest expression in nuclear and chromatin-bound fractions seen after TCPOBOP treatment in both male and female liver (**Fig. 6D; Fig. S6E, Fig. S6F**). Bright smFISH spots for both lnc5998 and Cyp2b10 appeared in many nuclei, indicating co-localization of the transcripts at the site of transcription.

**Fig. 6.**
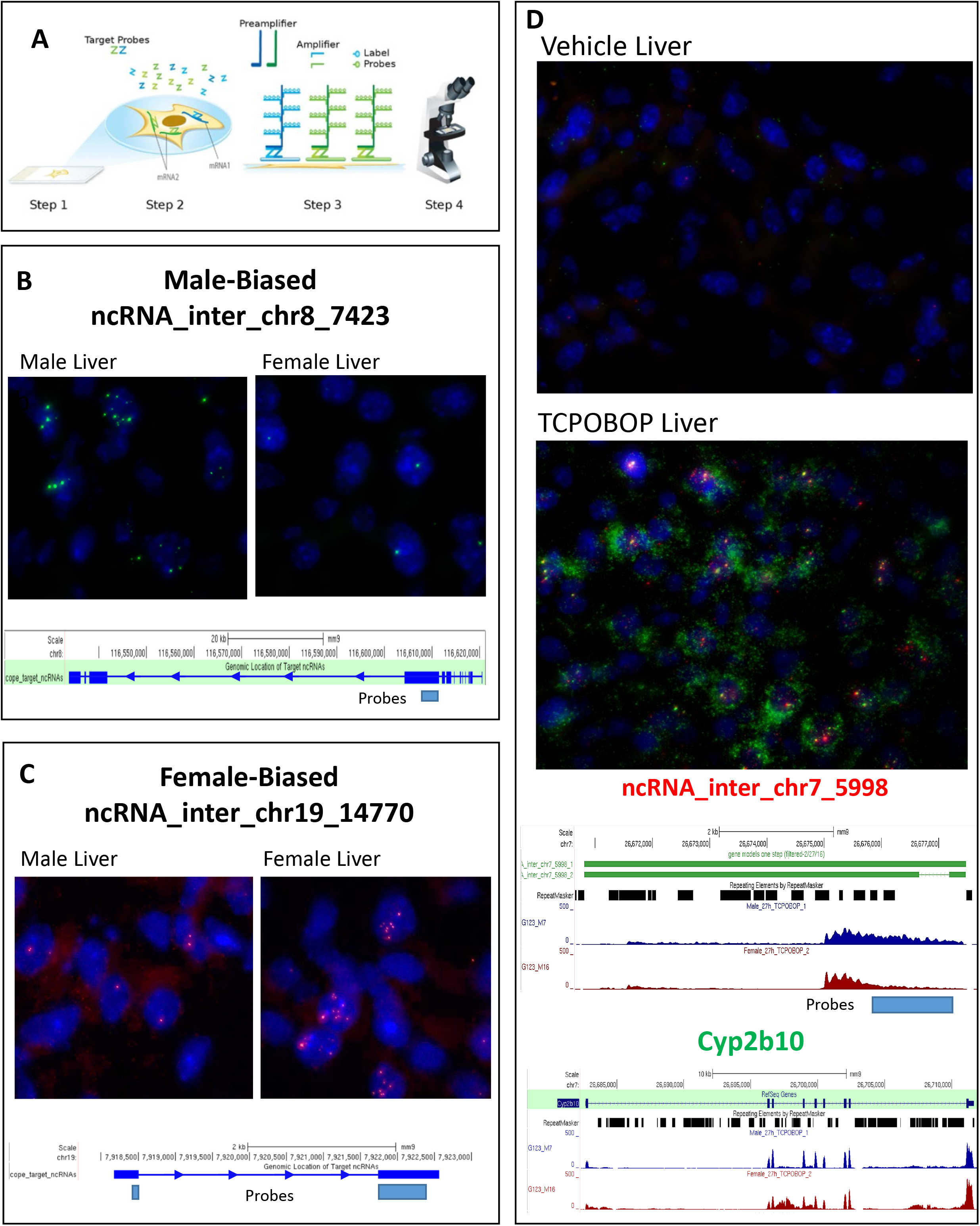
smFiSH analysis of sex-biased and TCPOBOP-responsive lncRNAs and PCGs. (**A**) Schematic from ACD, Inc describing how RNAScope technology works. ZZ probes are hybridized to the gene of interest, the pre-amplifier and amplifiers build in a tree-and-branch manner to amplify the probe/label signal up to 8000-fold, allowing for visualization of single RNA molecules. Visualization of expression for: (**B**) ncRNA_inter_chr8_7423, a male-biased lncRNA; (**C**) ncRNA_inter_chr19_14770, female-biased lncRNA, using ZZ probes designed to a small region of each target lncRNA; and (**D**) ncRNA_inter_chr7_5998, a TCPOBOP-inducible lncRNA, and the nearby PCG, Cyp2b10, using ZZ probes designed to a small region of lnc5998, and across the exonic structure of Cyp2b10. See Fig. S6 for quantitative analysis of smFISH data for all four genes.

### Integration with prior liver RNA-seq datasets

We integrated the above sets of regulated lncRNAs with prior, published datasets to help identify lncRNAs that are strong candidates for regulatory roles. We designated 49 lncRNAs as robust sex-biased based on their significant sex-biased expression in at least 2 of 5 subcellular fractions analyzed here (**Table S3A**) and in at least 5 of 11 prior liver RNA-seq datasets (**Table S4A**). These 49 lncRNAs are highly expressed and strongly sex biased: 40 show a maximum FPKM > 2, and 41 show a > 4-fold sex-bias in at least one of the five subcellular fractions. **Fig. 7A** presents expression data in both sexes across the subcellular fractions for eight of these lncRNAs, and highlights the large increases in expression, and hence the increased sensitivity for detection, seen in the chromatin-bound fractions. A large majority (86%) of the robust sex-biased lncRNAs showed a significant change in expression in livers of hypophysectomized mice, where the growth hormone signaling that regulates sex-biased gene expression is ablated [39]. Furthermore, 19 of the 49 lncRNAs exhibited developmental changes in expression in male mouse liver during the transition from the pre-pubertal stage to young adulthood [29], which has been linked to the sex-dependent expression of key transcription factors and sex-biased genes involved in specialized liver functions [40], suggesting these lncRNAs may contribute to the post-pubertal changes in expression commonly seen for sex-biased PCGs in male liver. Finally, 33 of the 375 sex-biased lncRNAs identified here showed significant sex-biased expression in two or fewer of the 11 prior datasets. A large majority (29/33) were maximally expressed in one of the chromatin-bound fractions, which helps explain why they were not detected previously by RNA-seq analysis of total liver or liver nuclear RNA.

**Fig. 7.**
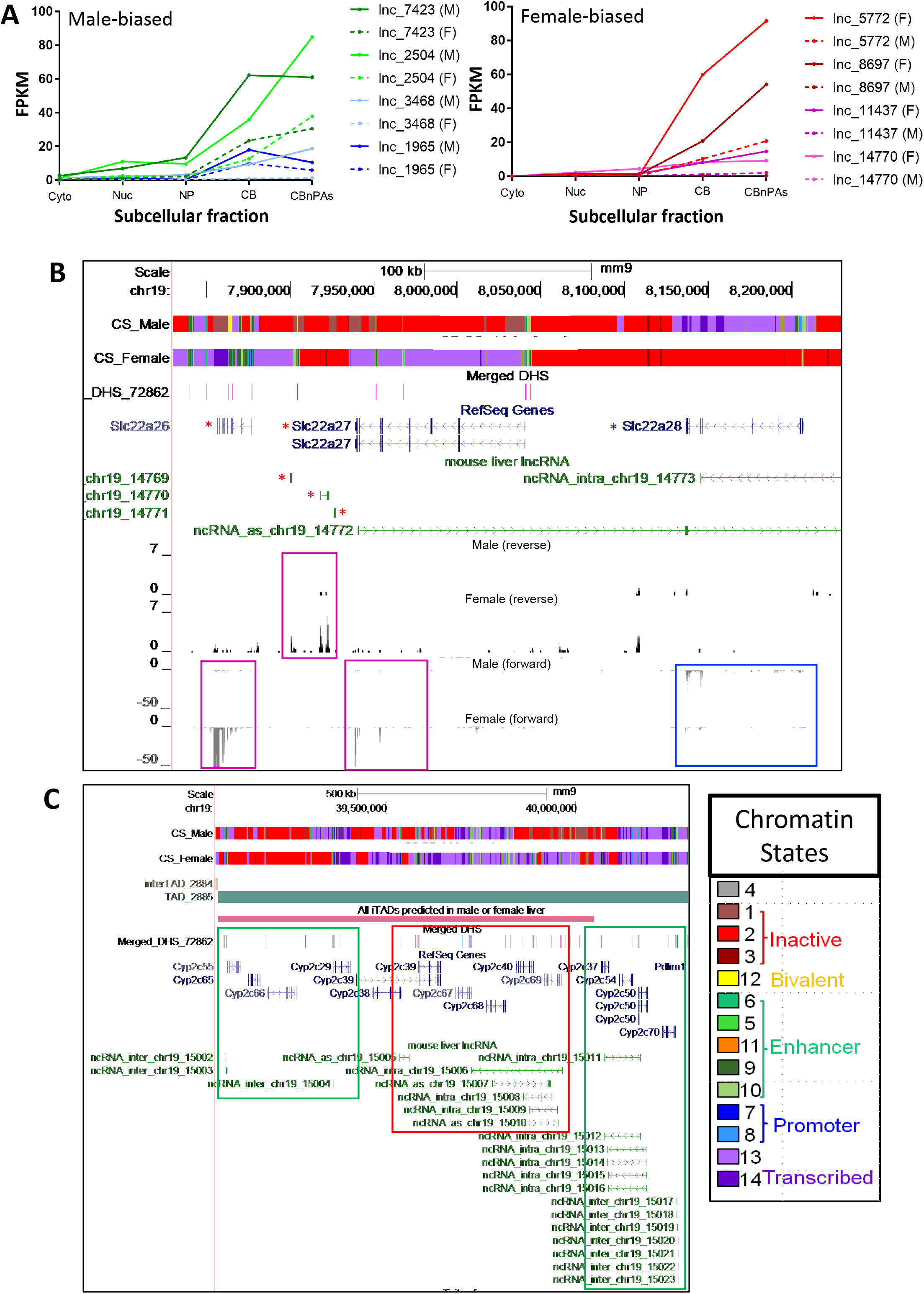
Sex-biased and TCPOBOP-responsive lncRNAs. (**A**) Robust female-biased (top) and male-biased lncRNAs (bottom) (Table S4A, column I) and their expression levels across five subcellular compartments (data based on Tables S2A-S2C). Solid lines indicate expression level in livers of the dominant sex, and dashed lines indicate expression in the opposite sex. Also see Table S3A. (**B**) and (**C**) UCSC genome browser screen shots highlighting individual sex-biased genes. The first two tracks in each panel describe the chromatin state in untreated male and female liver (red = inactive state, green = enhancer state, purple = actively transcribed state; see key of all 14 chromatin states at bottom right, based on {Sugathan, 2013 #321}). Also shown are tracks indicating the TAD structure (horizontal bars), where marked, followed by the Merged DHS track, which marks DHS that are female-biased (pink vertical bars), male-biased (blue bars) and sex-independent (gray bars). Tracks presenting RefSeq gene structures (blue) and lncRNA structures (green) are shown next. (**B**) Three female-biased lncRNAs and two female-biased PCGs (red asterisks, pink boxes), all within a TAD that also contains the male-biased gene Slc27a28 (blue box) (see Table S6A). Shown at the bottom are four tracks with normalized RNA-seq reads in the nucleoplasmic fraction for vehicle-treated male and female liver. on the forward and reverse strands, as indicated below each label. (**C**) TAD containing many Cyp2c genes and lncRNAs. The genomic region shown is divided into 3 regions whose genes are either up regulated (green boxes) or down regulated (red box) by TCPOBOP exposure (see Table S6B). The 4^th^ track from top shows a predicted intra-TAD (iTAD) loop (long red horizontal bar) that is found in female liver only.

We also designated 96 TCPOBOP-responsive lncRNAs as robust (responsive in at least 2 of 5 subcellular fractions, and in at least 2 of 5 prior TCPOBOP-treated datasets; **Table S5A**). These lncRNAs are highly expressed (79 have maximum FPKM > 2) and highly responsive to TCPOBOP (74 show > 4-fold maximum response). Many also responded to other chemicals that dysregulate gene expression in the liver, including phenobarbital (n=45 lncRNAs), acetaminophen (n=27) and agonists of PPARA (either WY14634 or fenofibrate) (n=33). 334 of the 1,005 TCPOBOP-responsive lncRNAs identified here responded in at most one of the five prior TCPOBOP-treated datasets. 70 of these 334 novel lncRNAs also responded to at least one of four other chemicals examined (phenobarbital, acetaminophen, WY14634 and fenofibrate), and 282 (84%) were maximally expressed in one of the chromatin-bound fractions, which may explain why they were not identified previously. The novel TCPOBOP-inducible lncRNAs include lnc4278/DANCR, lnc14777/SNHG1, lnc10895/SNHG10, which promote hepatocellular carcinoma through their actions as miRNA sponges [41-43], as well as lnc733/GAS5, which is also a miRNA sponge and acts to inhibit liver fibrosis [44].

### LncRNAs as potential regulators in *cis*

Many lncRNAs function as regulators in *cis*, whose transcription regulates the expression of nearby PCGs through a variety of mechanisms [8, 45, 46]. To identify sex-biased and TCPOBOP-responsive lncRNAs that may serve as *cis*-regulators, we considered lncRNAs co-localized with PCGs within TADs [47]. TADs are megabase-scale chromatin loops organized by interactions between cohesion and CTCF [48]. TADs loop together relatively distant regions of chromatin, allowing regulatory elements and their bound factors, including chromatin-tethered lncRNAs on one end of a TAD to regulate in cis genes located on the other end of the TAD. Using TAD definitions for mouse liver [48] (**Table S6C**), we identified 36 TADs that contain at least one strongly sex-biased lncRNA (>4-fold sex difference in expression) and harbor at least one sex-biased non-lncRNA gene (**Table S6A**). These 36 TADs encompass 93 sex-biased lncRNAs, 71 sex-biased PCGs and 4 sex-biased non-coding RefSeq genes which serve as candidates for *cis* regulation. 30 of the 93 lncRNAs are within 13 TADs that each contain at least one gene of the opposite sex bias, and hence is a candidate for negative regulation.

One example, is a genomic region defined by a gap between two TADs (inter-TAD region, 960 kb long) that contains three female-biased lncRNAs, two female-biased PCGs and one male-biased PCG (**Fig. 7B**, red and blue asterisks). The three PCGs are sex-biased in all five subcellular compartments: Slc22a26 and Slc22a27 show very strong female bias (>40-fold) and are highly expressed (FPKM = 63 and 7.9, respectively), while Slc22a28 is 4-fold male-biased with an FPKM (max) of 15 in the nucleoplasm. All three lncRNAs show strongly female-biased expression (14-fold to 66-fold in the subcellular fraction with the highest expression) with FPKM values as high as 4.5-7.5 in the nucleoplasm or in a chromatin fraction **(Tables S2A-S2C)**. The three lncRNAs are located between the two female-biased Slc22a genes in a region with multiple female-biased DNase hypersensitive sites (DHS), suggesting the entire TAD is regulated as one unit (**Fig. 7B**). In contrast, the region encompassing the male-biased Slc22a28, located > 100 kb upstream of the female-biased genomic region, is devoid of DHS sites; however, that genomic region is characterized by an active chromatin state in male but not female liver (**Fig. 7B**, top two tracks). Slc22a28 could be regulated by the one male-specific DHS found in the same TAD, 342 kb upstream of Slc22a28; alternatively, the female-biased lncRNA(s) could act via TAD-based looping to negatively regulate Slc22a28 and silence its expression in female liver, resulting in the observed male-biased expression.

We also identified 211 TADs that contain at least one TCPOBOP-responsive lncRNA (fold change > 4) and at least one TCPOBOP-responsive non-lncRNA, comprising a total of 484 lncRNAs, 418 PCGs and 28 non-coding RefSeq genes (**Table S7B**). One interesting example is a TAD that encompasses 6 TCPOBOP-regulated lncRNAs (4 induced, 2 repressed) and 13 TCPOBOP-regulated CYP2C gene subfamily PCGs (8 induced, 5 repressed) (**Fig. 7C**). This TAD encompasses 3 segments, the first and the third segment containing TCPOBOP-induced genes, and the middle segment containing TCPOBOP-repressed genes. This arrangement suggests that the TAD is divided into 3 insulated regions in TCPOBOP-exposed liver, suggesting there may be intra-TAD loops (i.e., sub-TAD structures) in this region. The first segment includes 5 up-regulated genes, including Cyp2c55, which is induced by TCPOBOP > 222-fold (to FPKM = 333). The strongest of the two induced lncRNAs in this segment, lnc15004, is induced 212-fold (FPKM = 46) and is a robust TCPOBOP-responsive lncRNA. The middle segment contains 4 lncRNAs and 5 PCGs, all of which are repressed by TCPOBOP. The highest expressed lncRNA (FPKM (untreated liver) = 7.2) is lnc15006, which is down-regulated 2.7-fold by TCPOBOP. Cyp2c69 (FPKM (untreated liver) = 187) is 12-fold down-regulation by TCPOBOP. The third segment of the TAD contains three PCGs and two lncRNAs that are up regulated by TCPOBOP. Cyp2c54 expression increases 30-fold (to FPKM = 1,671), while the robustly responsive lncRNA, lnc15014, is 80-fold induced (to FPKM = 33) **(Fig. 7C)**. Interestingly, there is evidence for an intra-TAD loop in untreated female but not male liver [49] that encompasses the first 2 segments, and excludes the third segment (**Fig. 7C**, 4th track, red horizontal bar). Of note, a majority of the genes in the TCPOBOP-repressed segment are more highly responsive to TCPOBOP in female liver, where 4 of 5 genes show female-biased expression in vehicle-treated liver. This female-specific intra-TAD loop could allow the robust induced lnc15004 to repress expression of these genes in TCPOBOP-treated female liver.

### Divergently transcribed lncRNAs showing sex-biased expression

Divergently lncRNAs are defined as lncRNAs with a TSS < 5 kb from the TSS of a non-overlapping PCG transcribed from the opposite strand, and are frequently adjacent to regulatory genes, whose expression or activity is controlled in cis by the divergently transcribed lncRNA [45, 50]. Accordingly, one can infer the biological function of a divergent lncRNA from that of its neighboring adjacent gene. We identified six sex-biased lncRNAs that are divergently transcribed from a sex-biased PCG (**Table S4A**, column AI). One of these divergent gene pairs involves SOCS2, a STAT5-induced inhibitor of STAT5 signaling [51, 52] that showed 3 to 5-fold female-biased expression (**Table S6A**). SOCS2 and other SOCS family proteins are negative feedback regulators of STAT5-dependent growth hormone signaling [52] and are proposed to contribute to the inhibitory effects of persistent growth hormone stimulation on STAT5 signaling in female liver [53]. SOCS2 also inhibits metastasis in hepatocellular carcinoma [54], a male-predominant disease [55, 56]. Lnc9183 is divergently transcribed from SOCS2 (**Fig. 8A**) and showed 9-fold female-biased expression, which was highest in non-polyA-selected chromatin-bound RNA (FPKM = 13; **Table S4A**). SOCS2 has several isoforms, and the major transcript in the cytoplasmic, nuclear and nucleoplasmic fractions has its TSS within a subTAD structure together with the TSS of its divergent sex-biased lncRNA partner. This genomic organization may insulate lncRNA-driven regulation of SOCS2 in female liver from other genes within the TAD that are not sex-biased. Of note, SOCS2 and the divergently transcribed lnc9183 are both TCPOBOP-responsive (**Table S6B**), as are several other, more distant lncRNAs in the same TAD (lnc9185 and lnc9178), which could impact STAT5 regulation of its many downstream sex-biased gene targets in male and female liver [57].

**Fig. 8.**
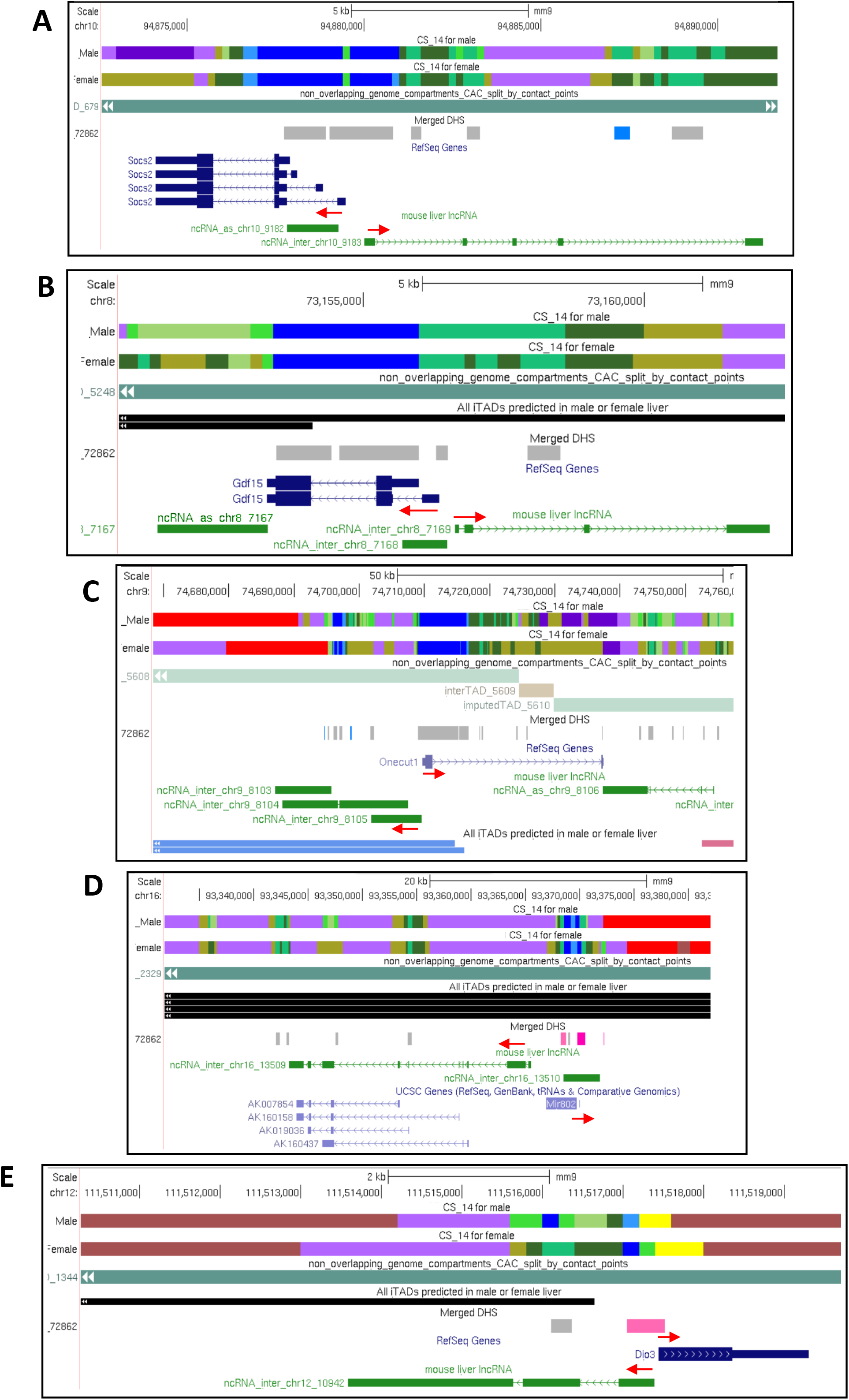
Divergently transcribed lncRNA-PCG genes of interest. Shown are UCSC genome browser screen shots, as described in Fig. 7, highlighting five divergent lncRNA-PCG gene pairs, whose direction of transcription is marked by red arrows. See data in Table S4A, column AI, and in Table S5A, column AW. (**A**) Divergent sex-biased and TCPOBOP-responsive gene pair, lnc9183 and Socs2. (**B**) Divergent TCPOBOP-responsive gene pair, lnc7169 and Gdf15. These genes fall within a sex-independent intra-TAD loop (black) with several sex-independent DHS (gray). (**C**) Divergent TCPOBOP-responsive gene pair, lnc8105 and Onecut1 (Hnf6), both of whose TSS are in the same male-biased intra-TAD loop (last track, light blue). (**D**) Divergent TCPOBOP-responsive gene pair, lnc13509 and Mir802. These genes fall within a sex-independent intra-TAD loop (black) with several female-biased DHS (pink) in the promoter regions of both genes. (**E**) Divergent TCPOBOP-responsive gene pair, lnc10942 and Dio3. These genes are excluded from the predicted intra-TAD loop nearby (black) and their promoters are both within a shared female-biased DHS region (pink).

### Divergent transcription of TCPOBOP-responsive lncRNA and PCG gene pairs

We also identified 51 divergently transcribed, TCPOBOP-responsive lncRNA-PCG pairs in either male or female liver (**Table S5A**, column AW). Four examples are described below.

**Lnc7169** is divergently transcribed from Gdf15 **(Fig. 8B)**, a stress response cytokine [58]. Lnc7169 has both rat and human orthologs, and both it and Gdf15 are strongly induced by TCPOBOP (5-42 fold and 16-18 fold, respectively; **Table S6B**). The rat ortholog maps to a gene network involving glycerolipid metabolism, and is dysregulated by 16 of 27 xenochemicals tested in rat liver, representing 6 of 7 xenochemical mechanisms of action [33]. Gdf15 impairs progression of non-alcoholic fatty liver disease in obese mice by enhancing fatty acids oxidation [59, 60] and its deficiency promotes high fat diet-induced obesity [61] and exacerbates liver injury induced by chronic alcohol and carbon tetrachloride exposure [62]. We hypothesize that the multi-xenobiotic-responsive lnc7169 plays a key role regulating these beneficial Gdf15-dependent hepatic disease-related processes.

**Lnc8105** is divergently transcribed from Onecut1/HNF6, a major liver-enriched transcription factor implicated in the expression of many liver-specific genes [22, 57, 63] **(Fig. 8C)**. TCPOBOP strongly down regulated both genes, in both male and female mouse liver. This repression of Onecut1/HNF6 RNA was observed in the chromatin fraction, indicating a decrease in of Onecut1/HNF6 transcription, but surprisingly, it did not result in a decrease in cytoplasmic Onecut1/HNF6 RNA (**Table S3B**). The impact of this repression by TCPOBOP of Onecut1/HNF6 and its divergently transcribed, chromatin-associated lncRNA on the many downstream transcriptional targets of HNF6 in mouse liver [63] are unknown.

**lnc13509 (**1810053b23Rik) is transcribed divergently from miR802 (**Fig. 8D**). Both genes are strongly induced by TCPOBOP (>15-fold; **Table S6B)**. Lnc13509 is highly expressed in fetal liver and then repressed during development [29]. It has a close human ortholog (57% identity) [33] and its rat ortholog is induced by multiple xenobiotics and is in a co-expression network of xenobiotic-responsive genes enriched in transcription factors [33]. We hypothesize that the induction of lnc13509 impacts various biological and pathological responses regulated by miR802. Elevated expression of miR802 is seen in type-II diabetes [64] and in livers of high fat diet-fed mice, where it impairs glucose metabolism by silencing the transcription factor HNF1b and by increasing oxidative stress [65-67]. Further, miR802 shows female-biased expression in mouse liver; it preferentially represses many male-biased mRNAs and increases levels of female-biased mRNAs in female liver [68] and has been associated with regulation of glucose and lipid metabolism [67].

Finally, **lnc10942/Dio3os** is repressed up to 20-fold in mouse liver by agonists of the nuclear receptor CAR but was induced 5-fold by PPARA agonists (**Table S5A, Table S5C**). Its rat ortholog is responsive to 13 out of 27 xenochemical exposures in rat liver [33]. Repression of Dio3os has been linked to increased cell proliferation, and its repression is a biomarker for inflammatory bowel disease [69, 70]. Lnc10942/Dio3os is divergently transcribed from deiodinase-3 (Dio3) (**Fig. 8E**), a seleno enzyme that inactivates the thyroid hormones T3 and T4 [71]. Systemic thyroid hormone inactivation occurs upon CAR activation in mouse liver, and is also a physiological response that limits weight loss upon fasting/caloric restriction [72, 73]. Further, Dio3 undergoes translational repression in models of drug-induced inflammation and hepatotoxicity. Given the opposite effects of CAR vs. PPARA activation on the expression of lnc10942/Dio3os, this lncRNA may contribute to some of the differing physiological effects of CAR vs PPARA activation in liver. Lnc10942/Dio3os may regulate responses to hepatic stressors, such as fasting, high fat diet, and the need to regulate thyroid hormone levels by Dio3.

## Discussion

Global patterns of gene expression, maturation and subcellular localization were determined for thousands of liver-expressed lncRNAs and PCGs using a fractionation protocol that allowed us to isolate, from the same individual mouse liver, cytoplasmic and nuclear RNA, as well as a soluble nucleoplasmic RNA fraction and RNA tightly bound to chromatin. Transcripts enriched in the chromatin-bound fraction were the least mature, as indicated by a high fraction of sequence reads mapping to introns, while cytoplasmic transcripts were the most mature. In contrast to PCGs, lncRNAs were highly enriched in the nucleus and specifically in the chromatin-bound fraction, rather than the nucleoplasm. Furthermore, many lncRNAs were most highly expressed in non-polyA-selected chromatin-bound RNA, consistent with findings in human cell lines that lncRNAs, as well as chromatin-enriched RNAs, are less poly-adenylated than mRNAs [28]. The increased sensitivity for lncRNA detection in chromatin-bound RNA enabled us to identify 375 lncRNAs showing sex-biased expression, as well as 1,005 lncRNAs that were significantly induced or repressed in livers from mice treated with the CAR agonist TCPOBOP, many of which were not identified in earlier work where nuclear but not chromatin-bound RNA was analyzed [21, 22, 29]. Finally, we identified lncRNAs associated with divergently transcribed lncRNA-PCG pairs, many of which are anticipated to have regulatory functions [50], as well as candidates for *cis*-acting lncRNAs [46, 74], based on their presence in the same TAD [47] as correspondingly responsive, or in some cases oppositely responsive PCGs.

We used normalized intronic to exonic read densities (IO/EC ratio) to assess the extent of transcript splicing in each cell fraction. This approach is similar to calculating the degree of splicing based on exonic base coverage divided by base coverage over the entire transcript [27]. We found splicing was comparatively low for both lncRNAs and PCGs in the chromatin-bound fraction, independent of polyadenylation selection. This supports the proposal that lncRNA and PCG splicing initially proceed in a similar manner, with polyA addition preceding, or occurring at the same time, as splicing [75]. However, while PCG transcripts became increasingly more mature in moving from chromatin to the nucleoplasm and then on to the cytoplasm, lncRNA transcripts showed less splicing than PCG transcripts in the nucleoplasmic, nuclear and cytoplasmic compartments, despite the presence of multiple splice sites junctions in many lncRNA transcripts. This finding is consistent with earlier work indicating that lncRNAs do not necessarily need to be spliced to be functional [36, 76].

Comparing transcript levels between subcellular compartments, we found that lncRNAs were highly enriched in the nuclear and chromatin-bound fractions and were substantially depleted from the cytoplasm and the nucleoplasm. Thus, at least 92% of liver lncRNAs showing significant differential expression between cytoplasm and nucleus were enriched in the nucleus, and 99% of lncRNAs differentially expressed between chromatin and nucleoplasm were chromatin-enriched. Chromatin retention of lncRNAs is thought to be mediated, at least in part, by inefficient splicing due to weak internal splicing signals and the associated increase in Pol II occupancy on introns compared to PCGs [77], which suppresses splicing-coupled mRNA export; however, we did not observe an association between lncRNA transcript maturity and enrichment for chromatin binding. Presumably, other features of lncRNAs, including the presence of specific *cis*-elements that mediate nuclear retention (e.g., repeat insertion domains, SINE-derived localization elements [78, 79] and motifs associated with U1 snRNP binding [80]) contribute to the tight chromatin binding that we observed. This chromatin binding is likely to be an important driver of many nuclear lncRNA functions, including direct or indirect regulation of chromatin states and gene transcription. Chromatin-bound lncRNAs may act in *cis* [46] at sites in the genome close to their transcription, but some may transit through the nucleoplasm and be *trans*-acting [8]. Of note, many lncRNAs enriched in the chromatin fraction were also present in the nucleoplasm at a significant level, which could allow them access to multiple *trans* sites within the nucleus.

The majority of the liver lncRNAs we examined appear to be poly-adenylated, insofar as they were recovered from polyA-selected RNA. However, by comparing a polyA-selected to a non-polyA-selected chromatin fraction, we found that a large fraction of chromatin-bound lncRNAs were enriched in non-polyA-selected RNA. Moreover, a distinct subset comprised of 506 chromatin-bound lncRNAs, as well as 26 PCGs, was apparently non-poly-adenylated, insofar as they showed >60-fold greater abundance in the non-polyA-selected fraction (Fig. 4B). Many chromatin-enriched lncRNAs are under-spliced compared to mRNAs [28] and yet appear to be functional despite incomplete splicing and/or poly-adenylation [36, 79]. Further, several subclasses of lncRNAs are not spliced, including very long intergenic lncRNAs, macro lncRNAs, and circular lncRNAs [76, 81]. Computational methods have been developed to predict lncRNA subcellular localization based on features such as splicing efficiency and the presence of certain k-mer sequences, specific binding motifs and genomic characteristics [77, 82]. Current methods are ∼ 75% accurate in predicting nuclear versus cytoplasmic localization of well characterized lncRNAs [82], and efforts at further refinement will benefit from experimentally validated datasets such as those described here.

PCG transcripts were more likely than lncRNAs to show both a cytoplasmic (vs. nuclear) bias and a nucleoplasmic (vs. chromatin-bound fraction) bias. Moreover, PCGs enriched in the cytoplasm were more extensively spliced in both the cytoplasm and the nucleus than their nuclear-enriched PCG counterparts. The same pattern was seen when comparing nucleoplasmic and chromatin-bound PCG transcripts, consistent with their localization bias largely being driven by transcript maturation. Indeed, chromatin-bound PCG transcripts showing a bias for the non-polyadenylated fraction were encoded by longer genes with a higher intronic content than transcripts biased toward the polyA-selected fraction; their enrichment in this fraction can thus be explained by the longer times required for gene transcription and splicing as compared to shorter, lower intronic content PCGs. mRNA export to the cytoplasm is facilitated by the completion of mRNA processing, and nuclear-retained mRNAs often contain introns [35, 83-85]. Nuclear retention of mRNAs can permanent, but may also be reversible in response to cell stressors [83]. These events are thought to aid in the stress response by stockpiling mRNAs for rapid release [86] and also to minimize fluctuations in protein levels due to bursty transcription [87].

Finally, we identified 375 lncRNAs showing sex-biased expression, as well as 1,005 lncRNAs responsive to the CAR agonist ligand TCPOBOP. Many of these lncRNAs were identified in chromatin-bound RNA, consistent with both these processes being regulated at the transcriptional level. The highest level of expression was often seen in the chromatin fraction, which increased the sensitivity for lncRNA detection and helps explain why many of these regulated lncRNAs were not discovered in earlier whole liver or total nuclear RNA-seq analyses. Many of these sex-biased and TCPOBOP-responsive lncRNAs may be *cis*-acting, based on their location together with similarly regulated, or in some cases oppositely regulated PCGs within the same TADs, where most promoter-enhancer interactions, including lncRNA-PCG interactions, are expected to occur [49]. Overall, 25% of sex-biased lncRNAs were located in TADs with other sex-biased genes. Similarly, 48% of TCPOBOP-responsive lncRNAs were in TADs with other TCPOBOP-responsive genes, giving them the potential to act in *cis*. We also identified 6 cases where sex-biased lncRNAs are divergently transcribed from correspondingly sex-biased PCGs, and 51 cases of divergently transcribed TCPOBOP-responsive lncRNA-RefSeq pairs. The presence of close orthologs in rat or human for several of the divergently transcribed, xenobiotic-responsive lncRNAs [33] supports their proposed functional roles in liver responses to foreign chemical exposure. In one example, the strong induction of lnc7169 by TCPOBOP may contribute to the hepatoprotective effects of CAR activation on high fat diet-induced non-alcoholic fatty liver disease [88, 89] by increasing expression of the divergently transcribed Gdf15, a stress response cytokine that is induced by inflammation, acute injury and oxidative stress [58]. In contrast, the very strong induction by TCPOBOP of lnc13509 may stimulate the divergent transcription of miR802, whose expression is elevated in type-II diabetes [64] and in livers of high fat diet-fed mice, where it impairs glucose metabolism and increases oxidative stress [65-67]. Alternatively, lnc13509 could serve as hepatoprotective miRNA sponge [90] that depletes miR802, which may be investigated using a variety of experimental and computational approaches [91, 92], including innovative knockout technologies [30] that may uncover its biological functions and gene targets in the liver.

### Conclusions

We characterized global patterns of expression, maturation and sub-cellular localization for the mouse liver transcriptome, including more than 15,000 lncRNAs, many of which showed tight binding to chromatin. Sequencing chromatin-bound RNA greatly increased the sensitivity for detecting lowly expressed lncRNAs and enabled us to discover and localize hundreds of novel regulated liver lncRNAs, including lncRNAs showing sex-biased expression or responsiveness to an agonist ligand of the nuclear receptor CAR. Integration of our findings with prior studies identified strong candidates for lncRNAs that may regulate a variety of hepatic functions based on their co-localization within topologically associating domains, or their transcription divergent or antisense to PCGs associated with pathways linked to hepatic physiology and disease.

## Methods

### Animal studies

All mouse work was carried out in compliance with procedures approved by the Boston University Institutional Animal Care and Use Committee. Male and female CD-1 mice (strain Crl:CD1(ICR)), between 7 and 8 weeks of age, were purchased from Charles River Laboratories (Wilmington, Massachusetts). TCPOBOP (1,4-bis(2-(3,5-dichloropyridyloxy))benzene) (purchased from Santa Cruz Biotechnology; Chem Cruze, Cat. #SC-203291) was dissolved in DMSO at 7.5 mg/ml, and then diluted 10-fold into corn oil, followed by intraperitoneal injection of 4 µl per gram body weight (final dose: 3 mg TCPOBOP and 4 µl of 10% DMSO in corn oil, per kg body weight) at 8 AM (Boston University Lab Animal Care Facility; lights on at 7:30 AM, lights off at 7:30 PM). Mice were euthanized 27 hr later, at 11 AM, by cervical dislocation under CO_2_. Liver samples used in this study were excised and flash frozen in liquid nitrogen then stored at −80° C, and were obtained from Dr. Hong Ma of this laboratory.

### Isolation of cytoplasmic and nuclear RNA

All pipette tips used were RNase-free and DNase-free (Cat. #76322 series, VWR). RNA was isolated from livers from n=4 mice from each of four treatment groups: vehicle-injected males, 27 h TCPOBOP-treated males, vehicle-injected females, and 27 h TCPOBOP-treated females (**Table S1A**). The following buffers were prepared fresh daily and kept on ice for up to 2 h: **Base Solution**, 10 mM Tris-Cl, pH 7.4, 146 mM NaCl, 1 mM CaCl_2_ 21 mM MgCl_2_; **Lysis Buffer**, Base Solution containing 0.1% Triton X-100 (Cat. #T8787, Sigma), with 80 U/mL Protector RNase Inhibitor (Cat # 3335402001, Roche) added just prior to use; **ST Nuclei Wash Buffer (ST Buffer)**, Base Solution containing 0.01% BSA (Cat. #SRE0036, Sigma), with 80 U/mL Protector RNAse Inhibitor added just prior to use; **BSA Wash Buffer**, 1X PBS containing 2% BSA and 0.02% Tween-20, with 80 U/mL Protector RNase Inhibitor added just prior to use. To minimize premature tissue thawing and RNA degradation, a ∼250 mg piece of each of four livers per group was placed on dry ice, cut into 2-3 smaller pieces, and stored in an Eppendorf tube on dry ice until ready for further processing. The combined frozen and pre-cut liver pieces were transferred to a 3 mL glass-on-glass dounce homogenizer on ice containing 1 mL of Lysis Buffer. Keeping the homogenizer on ice, each liver sample was dounced for 10 strokes with pestle A (loose fit) followed by ∼10 strokes with pestle B (tight fit) until the sample was fully homogenized. Homogenization was performed in under one minute, while avoiding foaming and splattering of the sample. ST Buffer (1 ml) was then added, pipetted up and down a few times to mix, and the sample was then passed through a 40 µm cell strainer (Cat. # 10199-655, VWR) into a 50 mL conical tube on ice. A second 1 ml of cold ST Buffer was used to rinse the homogenizer and pestles, passed through the same 40 µm cell strainer and then combined with the homogenized sample. The homogenizer and pestles were then washed with Milli-Q water three times before being used to process the next sample. Homogenized samples were kept on ice until samples from all liver groups were ready to proceed to the next step. Each homogenized sample was divided into two 1.5 ml Eppendorf tubes, which were centrifuged at 500 x g in a swinging bucket centrifuge (Dynac Centrifuge, Clay Adams) for 5 min at 4° C to pellet the lysed cells. To avoid damage to the nuclei, a swinging bucket rotor was used to minimize the shear forces generated using a fixed-angle rotor. The supernatant was removed from the pelleted nuclei, and 250 µl was placed in an Eppendorf tube on ice to extract cytoplasmic RNA. To prevent RNA degradation, 750 µl of Trizol LS reagent was added immediately to the cytoplasmic fraction, which was vortexed for a few seconds, then stored at − 20° C. The pelleted nuclei were gently resuspended in 1 ml BSA Wash Buffer while combining the material from both tubes into one sample, followed by centrifugation in a swinging bucket rotor at 500 x g for 5 min at 4° C. The supernatant was discarded and the pellet was resuspended in 1 mL BSA Wash Buffer then passed through a 20 µm cell strainer (Cat # 43-50020-03, PluriSelect). The strained sample was transferred into two LoBind 1.5 mL Eppendorf tubes (Cat # 022431021, Eppendorf) on ice: one tube with 333 µl was used to isolate nuclear RNA; 667 µl of Trizol LS reagent was immediately added to that tube, which was vortexed for a few seconds then stored at −20° C. A second tube with 667 µl was used to fractionate the nuclear RNA, as described below.

### Fractionation of nuclear RNA

This protocol was adapted from [93]. **Fractionation Buffer** (1% Triton X-100, 20 mM HEPES (pH 7.5), 300 mM NaCl, 2 M Urea [Cat #5505UA, LifeTech], 0.2 mM EDTA, 1 mM DTT, with 250 U/mL Protector RNase Inhibitor) was prepared fresh each day. The nuclei from the nuclei isolation step, above, were re-pelleted at 500 x g in a swinging bucket rotor for 5 min at 4° C, and the supernatant was discarded. The nuclear pellet was resuspended by gently pipetting in 200 µl Fractionation Buffer, followed by incubation on ice for 10 min then centrifugation at 3,000 x g for 2 min at 4° C. A 180 µl aliquot of the supernatant, corresponding to the nucleoplasmic (NP) fraction, was removed and placed in a clean Eppendorf tube on ice. Immediately, the total volume was brought to 250 µl with Milli-Q water; 750 µl Trizol LS was then added followed by vortexing for a few seconds and storage at −20 °C. The remaining supernatant (∼20 µl) was carefully removed without disturbing the pellet and discarded. The pellet (i.e., the chromatin-bound (CB) fraction) was gently washed twice with 100 µl Fractionation Buffer, taking care to not disturb the pellet, and then centrifuged at 3,000 x g for 2 min at 4° C. The chromatin was then incubated for 30 min at 37° C in 50 µl DNase I solution (1X DNase I buffer containing 0.2 U/µl DNase I [Cat #M6101, Promega] and 0.25 U/µl Protector RNase Inhibitor), with gentle mixing by pipetting every 10 min. The final volume was brought to 250 µl with Milli-Q water. 750 µl Trizol LS was then added, the sample was vortexed briefly and stored at −20 °C.

### RNA isolation from subcellular fractions using Trizol LS

Frozen samples containing cytoplasmic, nuclear, nucleoplasmic and chromatin-bound RNA suspended in Trizol LS were thawed on ice, and vortexed for 10 s to fully resuspend each sample. Chloroform (isoamyl alcohol free, 0.2 ml) was then added to each sample, followed by vigorous vortexing for 15 s. Each sample was allowed to sit for 2-3 min at room temperature and then spun at 12,000 x g for 15 min at 4° C. The clear upper, aqueous phase was carefully transferred to a new centrifuge tube, while being careful to not disturb the genomic DNA at the interface. Isopropanol (0.5 ml) was added to each sample followed by vortexing for 10 s. Glycogen (Cat. #AM9510, ThermoFisher) was then added to each sample (1 µg per 20 µl reaction). Samples were vortexed and then incubated for 10 min at room temperature. Samples were centrifuged at 12,000 x g for 20 min at 4° C and the supernatant was discarded. The RNA pellet was washed with 1 mL of 75% ethanol by vortexing, followed by centrifugation at 7,500 x g for 5 min at 4° C. The ethanol wash was removed and samples were air dried for 5-10 min. Final RNA pellets were resuspended in 20 µl Milli-Q water, quantified on a Qubit instrument using the Qubit RNA HS Assay (Cat. #Q32852, Invitrogen) and stored at −20° C.

### qPCR analysis

RNA (0.5 µg) purified from each of four different subcellular fractions, and without polyA selection, was treated with DNase I (Cat. #M6101, Promega) to remove DNA contamination. cDNA was then synthesized using High-Capacity cDNA Reverse Transcription Kit (Cat #4368814, Applied Biosystems). qPCR was performed using primers specific to the RNAs for mouse 18S, Cyp2b10, Xist, Neat1, Elovl3 and pre-Elovl3, designed using Primer Express and Primer3 software (http://bioinfo.ut.ee/primer3-0.4.0/) (see **Table S1A** for primer sequences). Quantitative real-time PCR was carried out on a CFX384 Touch Real-Time PCR Detection System (Bio-Rad) using Power SYBR Green PCR Master Mix (ThermoFisher). Normalized linear Ct numbers were computed to determine the relative expression level of each gene across treatments and subcellular fractions to validate the effectiveness of RNA fractionation prior to sequencing library preparation.

### Single molecule imaging of RNAs in liver slices

Single molecule fluorescent in situ hybridization (smFiSH) to frozen mouse liver tissue slices was used to localize individual RNAs. We used RNAScope technology (Advanced Cell Diagnostics, Inc, Newark, CA) [38], which employs a series of up to twenty “ZZ” pairs of 20-mer oligonucleotides hybridized to each RNA transcript as a base for tree-and-branch building, leading to an overall 8,000-fold amplification of signal and enabling highly sensitive imaging and localization of single RNA molecules. ZZ probes were designed in cooperation with Advanced Cell Diagnostics staff for three lncRNAs (mouse mm9 genomic coordinates indicated): (1) 20 probes for lnc7423 across 2 exons, at Chr8(-):116,609,566-116,610,245 (680 bp), and at Chr8(-):116,609,152-116,609,565 (414 bp); (2) 9 probes for lnc14770 across 2 exons, at Chr19(+):7,918,477-7,918,504 (27 bp) and at Chr19(+):7,921,742-7,922,550 (808 bp); and (3) 20 probes for lnc5998 over 1 exon, where the majority of the expression is observed, at Chr8(-):26,676,223-26,677,271 (1,048 bp). We were unable to design a set of ZZ probes unique for Cyp2b10 due to its high homology with four other mouse Cyp2b subfamily members (Cyp2b9, Cyp2b13, Cyp2b19 and Cyp2b23). However, we did identify four ZZ probes spread across the exonic structure of Cyp2b10 that showed low homology with Cyp2b9 and Cyp2b13 (both liver expressed) but were homologous to Cyp2b19 and Cyp2b23, which are not expressed in mouse liver, and hence gave Cyp2b10-specific signals. The high specificity of these four ZZ probes for Cyp2b10 visualization was verified by the very low smFiSH signal in untreated male liver, where Cyp2b10 expression is very low. All other ZZ probes were unique to both coding and non-coding regions of the mouse genome.

Fresh mouse liver tissue was frozen in isopentane on dry ice, followed by flash freezing in liquid N_2_ and storage at −80° C. Frozen tissue was embedded in OCT medium and stored long-term at −80° C. Before cutting tissue slices, frozen tissue was allowed to incubate for 1 h at −20° C in the cooling chamber of Leica CM1950 cryostat. Tissue sections were sliced 15 µm thin and placed on SuperFrost Plus slides (Cat #12-550-15, Fisherbrand) and stored at −20° C. Slides were processed using RNAscope Fluorescent Multiplex Reagent Kit v1 for Fresh Frozen Tissue, as described in Advanced Cell Diagnostics documents #320513 and #320293, which lists all specialized reagents and equipment used for this protocol, except for 32% paraformaldehyde (Cat #15714-SP, Electron Microscope Science) and Prolong Gold Antifade with DAPI (Cat #8961, Cell Signaling Technology). Fixative (200 mL of fresh 4% paraformaldehyde in 1x PBS) was pre-chilled to 4° C. Groups of up to 8 slides with liver sections were taken directly from storage at −20° C and placed on a slide rack in pre-chilled fixative for 15 min at 4° C. Slides were then removed from the fixative and dehydrated by immersion in 200 mL of 50% ethanol for 5 min, followed by 5 min in 70% ethanol, and then twice in 100% ethanol for 5 min. Slides were incubated at −20° C in fresh 100% ethanol overnight. At the start of the next day, a water bath and an Advanced Cell Diagnostics HybEZ oven were set to 40° C after placing Advanced Cell Diagnostics humidity paper soaked in distilled water in the oven’s humidity control tray. Dehydrated slides were air dried on absorbent paper for 5 min at room temperature, and a hydrophobic barrier was drawn around each tissue slice using a special Advanced Cell Diagnostics marker pen and allowed to dry for 1 min. Slides were then placed on the HybEZ slide rack. Five drops of Pretreat 4 protease reagent was added to each section and then incubated for 30 min at room temperature with the incubation tray cover on. Materials for probe hybridization were prepared during this incubation step. 50x Wash Buffer (60 mL) was pre-warmed at 40° C for 10-20 min before adding to 2.94 L of Milli-Q water in a sealable container to make 1x Wash Buffer (stable at room temperature for over 1 month). Probes were warmed at 40° C for 10 min and cooled to room temperature before use, and the amplifying reagents (Amp1-FL to Amp4-FL) were warmed to room temperature. After 30 min, excess liquid was flicked off each slide and the slides were washed in 1x PBS twice by submerging the rack in the PBS wash 3-5 times. Slides were allowed to sit in 1x PBS for up to 15 min before proceeding to probe hybridization. Slides were then tapped gently to remove any excess liquid and placed on the HybEZ rack. Probe hybridization was performed using one of the following: mixture of up ZZ probe sets for up to three RNAs of interest, each using a different fluorescent channel; RNAscope 3-plex Positive Control probes (Cat # 320881); or RNAscope 3-plex Negative Control Probe (Cat # 320871). Four drops of the above described probe mixture was added to each slide and the rack was placed in the oven for 2 h at 40° C. After 2 h, the tray was removed from the oven, excess liquid was flicked from the slide, and the slide was washed twice in 1X wash buffer at room temperature for 2 min. This hybridization and washing procedure was repeated for each of four sequential amplification probes (Amp1-FL through Amp4-FL), with differing periods of incubation for each step: Amp1-FL for 30 min, Amp2-FL for 15 min, Amp3-FL for 30 min, and Amp4-FL for 15 min. We used the manufacturer’s Amp4 Alt A set-up (Cat. # 320855) for this experiment, which labels the Channel 1 probe with Alexa488 (green) and the Channel 2 probe with Atto 550 (orange). Due to the high auto-fluorescence of liver tissue in the green spectrum, we visualized the most highly expressed RNAs on Channel 1 and the less highly expressed RNAs on Channel 2. Thus, for the sex-biased lncRNAs, lnc7423 was visualized on Channel 1 and lnc14770 on Channel 2; for the TCPOBOP-inducible genes, Cyp2b10 was visualized on Channel 1 and lnc5998 on Channel 2. After the final wash step, excess liquid was removed by gently tapping, and 20 µl of Prolong Gold Antifade with DAPI (Cat #8961, Cell Signaling Technology) was placed in the center of the slide. A coverslip was added and the slide stored in the dark at 4° C.

Slides were imaged on a spinning disk confocal microscope (Olympus BX61) with a 60x oil immersion lens using preset channels to visualize DAPI (358 nm excitation/461 nm emission; blue), Alexa 488 (488/540 nm; green), Atto 550 (550/576 nm; orange) and Atto 647 (647/669 nm; far red). For processing, images were imported into FIJI image analysis software (https://imagej.net/Downloads) where channels were assigned to the proper visualization color and the z-stack was collapsed. Individual channels were separated and the delete background function was applied to each channel individually to remove excess noise, using a rolling ball radius of 50 pixels. Single channels were then merged to create a final image that was saved in RBG format to preserve the settings and converted to a .tif file. To quantify signal, the count dots feature of FIJI image analysis was applied to each channel individually, excluding DAPI, and counts were normalized over 5 fields of view for each measurement. Nuclei were counted manually for each image due to the variation of DAPI staining in the nucleus caused by euchromatin and heterochromatin.

### RNA-seq library preparation and sequencing

Sequencing libraries were prepared for 13 validated mouse livers (3 vehicle-treated males, 3 TCPOBOP-treated males, 3 vehicle-treated females, and 4 TCPOBOP-treated females) using RNA purified from each of the four subcellular fractions described above (cytoplasm, nucleus, nucleoplasm, chromatin-bound fraction). Illumina sequencing libraries were prepared using 0.5 µg of input RNA by poly(A) selection using the NEBNext Poly(A) mRNA Magnetic Isolation Module (Cat #E7490L), followed by library synthesis using the NEBNext Ultra Directional RNA Sequencing for Illumina kit (Cat #E7420L). An additional 0.5 µg of each of the 13 chromatin-bound RNA samples was also processed without poly(A) selection (CBnPAs fraction) to give a total of 13 livers x 5 fractions each = 65 RNA-seq libraries. Illumina sequencing was carried out by Novogene, Inc (Sacramento, CA) and yielded a total of 1.68 billion 150 bp paired-end read sequence fragments. Raw and processed sequencing flies are available for download from GEO (https://www.ncbi.nlm.nih.gov/geo/) accession <u>GSE160722</u>. Counting and mapping statistics for each sequenced sample are found in **Table S1A**.

### Sequence read counting

RNA-seq data was processed using a custom pipeline described elsewhere [31]. Custom Gene transfer format (GTF) files previously used to count sequence reads mapping to 24,197 RefSeq genes and 15,558 liver-expressed lncRNA genes, which are numbered sequentially by genomic coordinates [22], were combined into single GTF files, after removing 851 non-coding RefSeq genes that significantly matched one of the 15,558 lncRNA structures (608 identical matches plus 243 partially overlapping genes; see below) to avoid duplicate counting of those genes (listed in **Table S1D**, column K). The extent of match was initially determined by Bedtools intersect and Bedtools coverage commands, with > 30% overlap of the lncRNA structure with a non-coding RefSeq gene deemed to be a significant match. Overlaps ≤ 30% were then manually curated to identify RefSeq-lncRNA pairs with highly similar exonic features, as determined by visual assessment and best judgement. Based on these analyses: 608 RefSeq non-coding RNAs showed near perfect overlap (>98% match) with our set of 15,558 lncRNA structures; 319 RefSeq genes (most of which were non-coding genes) were 98% contained within the longer lncRNA structure; and 62 lncRNAs were >98% within a longer RefSeq gene, most of which were protein-coding RNAs. Of the 319 RefSeq genes found within longer lncRNAs, 243 were determined to be the same gene, based on the criteria described above, and 1 of the 62 RefSeq genes encompassing a shorter lncRNA was considered to be the same gene. Two genes were lost due to Excel errors generating double entries for gene name converted to 1-Mar and 2-Mar (https://www.theverge.com/2020/8/6/21355674/human-genes-rename-microsoft-excel-misreading-dates), leading to the final total of 38,901 RefSeq + lncRNA genes.

Three separate GTF files were prepared for the set of 38,901 genes (**Supplemental Files S1, S2, S3**), comprised of the following features for each gene: (1) Gene Body GTF, which includes the full genomic region of each gene, from the transcription start site to the transcript end site; (2) Exon Collapsed GTF, which includes all genomic regions that are exonic in any isoform of a gene; and (3) Intronic Only GTF, which includes all genomic sequences within intronic regions that are shared across all isoforms of a gene, i.e., regions do not overlap any exon in any isoform. Sequence reads were mapped using TopHat (v2.0.13), and multi-mapped reads were removed from the BAM files, leaving only singly mapped reads, which were counted by featureCounts (1.4.6-p5) using the above custom GTF files. The MultiOverlap option of featureCounts was enabled using the –O option, so that reads that overlap two or more genes in a GTF file were included in the counts for each gene. For many intragenic lncRNAs, the Gene Body counts and the Intronic Only counts were artificially high due to the inclusion of exonic reads from highly expressed, overlapping RefSeq protein-coding genes located within the lncRNA’s intronic regions. Similarly, many miRNAs are found within introns of highly expressed protein-coding genes, and consequently, their Gene Body counts are inflated due to the inclusion of spliced reads from the overlapping protein-coding genes. To mitigate these issues, the Gene Body and Intronic Only count files output by featureCounts were modified for all 249 intragenic lncRNA genes and all 1107 miRNA genes, as follows: Gene Body read counts were replaced by Exon Collapsed read counts, and Intronic Only read counts were set to zero. For those genes, the gene lengths used to calculate FPKM (fragments per kilobase length per million mapped sequence reads) values in downstream analyses were correspondingly modified to reflect the changes in counting regions for those genes.

In principle, the sum of read counts mapping to Exon Collapsed (EC) regions plus those that map to Intronic Only (IO) regions should equal the Gene Body (GB) read counts. However, for many genes, we found that the EC + IO counts exceeded the GB counts due to unspliced reads that cross an exon/intron boundary, which were included in both the EC counts and the IO counts. For most genes, the excess in EC + IO read counts over GB read counts was considered to have little impact, defined as either a difference of < 20% of the total read counts or a difference of fewer than 15 reads. Sequence read counts for EC + IO regions of genes that exceeded the corresponding GB region counts by at least 15 reads (average) were defined as either Minor (between 20% and 2-fold difference) or Major (>2-fold difference), with a 2-fold difference expected if all reads cross an intron/exon boundary are therefore double counted. Minor differences were uncommon, except in the chromatin-bound fractions, which contain many immature, unspliced transcripts (**Table S1B**). Major differences between EC + IO and GB reads were rare, being found for a total of only 9 RefSeq genes and 0 lncRNAs across all five fractions out of the 38,901 genes analyzed (**Table S1B, Table S1C**). In some case, GB read counts exceeded EC + IO read counts (resulting in Minor differences for 72 genes and Major differences for 54 genes, with the fewest such differences in the chromatin-bound samples). ∼82% of the genes GB > EC + IO read counts were antisense lncRNAs whose reads almost exclusively correspond to intronic regions of the opposite strand lncRNAs. Data for genes showing significant differences in read counts between EC + IO versus GB counting are summarized in **Table S1B** and **Table S1C**, and individual genes are flagged in the expression analysis summarized in **Table S1D**.

### Intronic/Exonic read density ratio

Genes with an intronic length of zero, i.e., all mono-exonic genes, were excluded from this analysis, as were all intragenic lncRNA and miRNA gene structures, due to the modifications to their counting described above. All other genes were separated into three categories: (1) Not Expressed, genes with an average across untreated male or female liver samples (n=3 each) of < 3 reads per sample in EC regions and also in IO regions; (2) Low Expressed, genes that have an average maximum across untreated male or female samples of 3 to 9 reads per sample in either EC or IO regions; and (3) Expressed, genes with > 9 reads per sample, averaged across untreated male or female samples, in either EC or IO regions. A total of ∼220-420 antisense lncRNAs, ∼360-780 intergenic lncRNAs and 11,700-13,700 RefSeq genes met the criteria for Expressed in either male or female liver each cellular compartment (**Table S1D, Table S1E**). RefSeq gene accession numbers and gene names were used to classify genes as protein-coding (NM accession numbers only), non-coding (NR accession numbers only), protein-coding/non-coding (both NM and NR accession numbers are assigned to different isoforms of the same gene), snRNAs and miRNAs, and to remove genes with NR accession numbers, which reduced the overall list of 24,197 RefSeq genes to a list of 20,082 RefSeq PCGs. In total, 1,442 multi-exonic lncRNA genes and 13,737 multi-exonic PCGs (**Table S1F**) were considered for this analysis across all 5 subcellular fractions, as presented below in **Fig. 2** for male liver and in **Fig. S2** for female liver; however, not all genes were expressed at a high enough level to be assessed for Intronic/Exonic read density in every fraction.

To calculate Intronic/Exonic read densities, we first added a pseudo-count of 0.1 reads to both the Exon Collapsed and Intronic Only read counts for each gene. Exon Collapsed and Intronic Only read counts for each gene were then normalized by the sequencing read depth of each sample, and the resulting normalized counts were averaged together across the n=3 biological replicates for both counting regions. To compare sequence read density in intronic versus exonic regions, we first computed for each gene the fraction of the full length gene body that is in an IO region, and the fraction that is in an EC region. The mean normalized read counts for IO and for EC regions were then divided by their respective fraction of full gene body length to give a genomic length-weighted normalized average read count. For each gene, the intron/exon read density ratio was then determined by dividing the weighted normalized average read counts for the IO region by that of the EC region (**Table S1F**).

### Differential expression analysis: subcellular compartmental bias

The pre-ribosomal RNA gene RNA45S was substantially removed during polyA selection and thus comprised only 0.2-14% of sequence reads in all except for the chromatin-bound non-PolyA-selected sequencing libraries, where it comprised 46-51% of all EC region reads; hence, RN45S sequence reads were removed from all 65 RNA-seq samples when analyzing differential expression between subcellular fractions. All samples were then normalized to 10 million EC region reads per sample, in order to compare the relative number of gene transcripts across the five subcellular fractions by differential expression analysis using edgeR (exact test). Differential expression comparisons were carried out between three pairs of subcellular compartments: (1) Cytoplasm vs Nucleus (both polyA-selected), (2) Nucleoplasm vs Chromatin Bound (both polyA-selected), and (3) Chromatin Bound (polyA-selected) vs Chromatin Bound (non-PolyA-selected). Each comparison was carried out for four biological conditions: untreated male and female liver, and separately, for 27-h TCPOBOP-stimulated male and female liver. **Tables S2A-S2C** present the differential expression results across all 4 biological conditions for each of the 3 pairs of subcellular fraction comparisons. For the final analysis of differentially expressed genes, only lncRNAs and PCGs were considered. A gene was considered to show a significant compartment bias if its differential expression between subcellular fractions met the stringent threshold of edgeR-adjusted p-value < 0.001 in at least one of the four biological conditions. Where indicated, analyses considered genes showing a compartment bias at the relaxed significance of edgeR-adjusted p-value < 0.05. The magnitude of the compartment bias was taken as the normalized ratio of the FPKM expression values for each subcellular fraction, determined using exon collapsed reads, and is presented for the biological condition with the highest FPKM value that shows significant bias. Graphs showing Subcellular fraction (Compartment) bias vs Fold Change for each of the three pairwise comparisons are presented below in **Fig. 3** and **Fig. 4**, with additional analyses in **Fig. S3, Fig. S4**, and **Fig. S5**.

### Discovery of sex-biased and TCPOBOP-responsive genes

Differential expression analysis was performed with edgeR using the exon collapsed read counts and gene length GTF definitions for each of the five subcellular compartments examined. The percentage of sequence reads derived from RN45S was consistent within each subcellular fraction for all four biological conditions, and so RN45S sequence reads were not removed for these differential expression analyses. Comparisons were carried out between three sets of biological conditions: (1) Untreated Male vs Untreated Female liver, (2) Untreated Male vs 27hr TCPOBOP-treated Male liver, and (3) Untreated Female vs 27hr TCPOBOP-treated Female liver. These comparisons were performed separately using sequence reads from each of the five subcellular fractions: cytoplasm, nucleus, nucleoplasm, chromatin-bound (all polyA-selected) and chromatin-bound non-PolyA-selected. Data on the sex-bias and TCPOBOP-responsiveness of all 38,901 genes across the five fractions is shown in **Table S3A** and **Table S3B**, respectively. For these analyses, a gene was considered sex-biased or TCPOBOP-responsive if it met an edgeR-adjusted p-value cutoff of 0.05 for differential expression in at least one of the five fractions. Sex-biased and TCPOBOP-responsive genes comparing fold-change to expression values (FPKM) in the significantly biased subcellular fraction with the largest FPKM are presented below in **Fig. 5A** and **Fig. 5B**, respectively.

### Integration of prior datasets: liver sex-differences

Prior published RNA-seq datasets comparing gene expression in untreated male vs untreated female mouse liver were integrated and compared with the results obtained in this study (**Table S4A, Table S4C, Table S4D**). Differential sex-biased expression, and responsiveness to hypophysectomy [29] or to deletion of Ezh1/Ezh2 [94] were determined at a threshold of > 2-fold expression difference at an edgeR-adjusted p-value (FDR) < 0.05. The prior RNA-seq datasets used in this analysis used either total, nuclear or cytoplasmic liver RNA, with either polyA selection or Ribo-minus ribosomal RNA depletion, and either CD-1 mouse livers [22, 31, 32, 95] or C57BL/6J mouse livers [96], as indicated in **Table S4**. Antisense lncRNAs were not considered for the datasets obtained by unstranded RNA sequencing, as the genomic strand of the sequence reads could not be determined. RNA-seq datasets comparing liver expression in intact male or intact female mice to that in hypophysectomized male or female mice were included to determine the response of sex-biased lncRNAs to loss of pituitary-dependent growth hormone signaling. Class 1 lncRNAs are those that, following hypophysectomy, show decreased expression in the sex where the lncRNA is more highly expressed in intact mouse liver. In contrast, Class 2 lncRNAs increase in expression following hypophysectomy in the sex where they show lower expression in intact mice [29]. Hypophysectomy class assignments were made for lncRNAs that showed sex-biased expression in at least one of the five subcellular fractions in this study, or in at least one of the prior datasets described above (**Table S4A**, column K). Sex-biased lncRNAs whose expression levels significantly change in male liver after 20 days of age, as we determined elsewhere [29], were assigned to four classes (**Table S4A**, column L). Interesting sets of robust, novel and non-responsive sex-biased lncRNAs (**Table S4A**, column I) are identified in **Table S4B**.

### Integration of prior datasets: TCPOBOP-responsiveness

The current analysis of genes showing TCPOBOP responsiveness in one or more subcellular fractions (**Table S3B**) was integrated with five prior datasets examining responsiveness to TCPOBOP (**Table S5C**): four polyA-selected nuclear RNA datasets from 3-h and 27-h TCPOBOP-exposed Male and Female liver vs sex-matched vehicle controls (series G123), and one polyA-selected total RNA dataset from 3-h TCPOBOP-treated Male vs vehicle-treated Male liver (series G95) [21]. LncRNAs were identified as significantly up regulated or significantly down regulated by TCPOBOP or other xenobiotic exposures (**Table S5C**) using a threshold of normalized absolute fold-change > 2 and an edgeR-adjusted p-value < 0.05. If datasets conflicted, e.g., where a gene was UP in one data set and DOWN in another, the response was characterized as MIXED. Other xenobiotic exposures examined (**Table S5C**) include: phenobarbital treatment of male liver (GSE77729), acetaminophen exposure of male liver (G111828), TCPOBOP or CITGO exposure of livers of mouse CAR (constitutive androstane receptor) mice or human CAR transgenic mice (GSE98666), and WY14634 or fenofibrate treatment of both mouse PPAR mice and human PPAR transgenic mice (series G134). Results are integrated in **Table S5A**, and descriptions of interesting groups of robust, novel and non-responding TCPOBOP-responsive lncRNAs (**Table S5A**, column I) are presented in **Table S5B**.

### Analysis of lncRNAs based on proximity and genomic organization

The RefSeq gene closest in linear distance to the gene body of each lncRNA gene, without regard to direction or genomic strand, was determined using the bedtools closest function. RefSeq genes that overlap a lncRNA gene on either strand were assigned a distance of zero. If multiple genes were discovered at the same distance away, all were considered in this analysis. Divergently transcribed RefSeq genes met all three of these criteria: transcription start site (TSS) within 5 kb of a lncRNA TSS; does not overlap the lncRNA gene; and is transcribed from the opposite strand. Genes that overlap a lncRNA gene were characterized as antisense if they are on the opposite strand; they were designated intragenic if they are on the same strand. The union of the Closest, Antisense, Intergenic and Divergent RefSeq gene lists was compared to six lists of RefSeq genes relating to liver function and disease: 326 sex-biased RefSeq genes (padj < 0.05) and 1,815 TCPOBOP-responsive RefSeq genes (padj < 0.05), both from this study across any subcellular fraction; and 772 non-alcoholic steatohepatitis-responsive genes based on single cell RNA-seq analysis [97], 107 non-alcoholic fatty liver disease-related genes [98], 217 liver-related fibrosis genes [94], and 920 hepatocellular carcinoma-related genes [94]. These gene lists are integrated in **Table S4A** and **Table S5A** (last columns). RefSeq genes that are within the same topologically associated domain (TAD) as lncRNAs showing sex-biased expression or TCPOBOP responsiveness in one or more subcellular fraction are shown in **Table S6**. Determination of TAD regions, including inter-TAD and imputed TAD region boundaries, was based on analyses performed by Gracia Bonilla of this laboratory, and was computed based on 3,538 TAD regions obtained from Hi-C data [48, 99] and 2,403 inter-TAD regions predicted computationally in male mouse liver [48] (**Table S6C**).

### Graphing and statistics

Error bars shown in scatter dot plot column graphs represent the inter-quartile range (IQR) of the distribution from the median. Statistical analysis for these graphs was performed using the Kruskal-Wasllis test followed by Dunn’s multiple comparison tests to obtain adjusted p-values, as implemented in the graphing software Prism to compare the distributions of multiple unmatched groups in a nonparametric manner.

## Ethics approval

All mouse work was carried out in compliance with procedures approved by the Boston University Institutional Animal Care and Use Committee, protocol PROTO201800698.

## Availability of data

The datasets generated and/or analyzed during the current study are included in this published article and its supplementary information files. Raw and processed sequencing files are available from GEO (https://www.ncbi.nlm.nih.gov/geo/) accession <u>GSE160722.</u>

## Authors’ contributions

CNG and DJW jointly conceived and designed the study. CNG carried out all of the laboratory experiments and a majority of the data analyses and prepared the figures and tables for publication. CNG and DJW jointly drafted the manuscript and DJW edited the manuscript for publication. Both authors read and approved the final manuscript.

## Acknowledgements

The authors thank Dr. Hong Ma for initiating nuclear fraction studies, and for providing the frozen mouse liver tissue used in this study, and Kritika Karri for her guidance in implementing the RNA-seq read counting methods utilized in these analyses.

## List of Abbreviations

CAR: constitutive androstane receptor
CBnPAs: chromatin-bound, non-polyA-selected
DHS: DNase hypersensitive site(s)
EC: exon collapsed
FDR: false discovery rate
FPKM: fragments per kilobase; length per million mapped sequence reads
GB: gene body
GTF: gene transfer format
IO: intronic only
IQR: inter-quartile range
lncRNA: long non-coding RNA
PCGs: protein coding genes
smFISH: single molecule RNA fluorescence in situ hybridization
TAD: topologically associated domain
TCPOBOP: 1,4-bis(2-(3,5-dichloropyridyloxy))benzene)
TSS: transcription start site

## Supplemental Figures

**Fig. S1.**
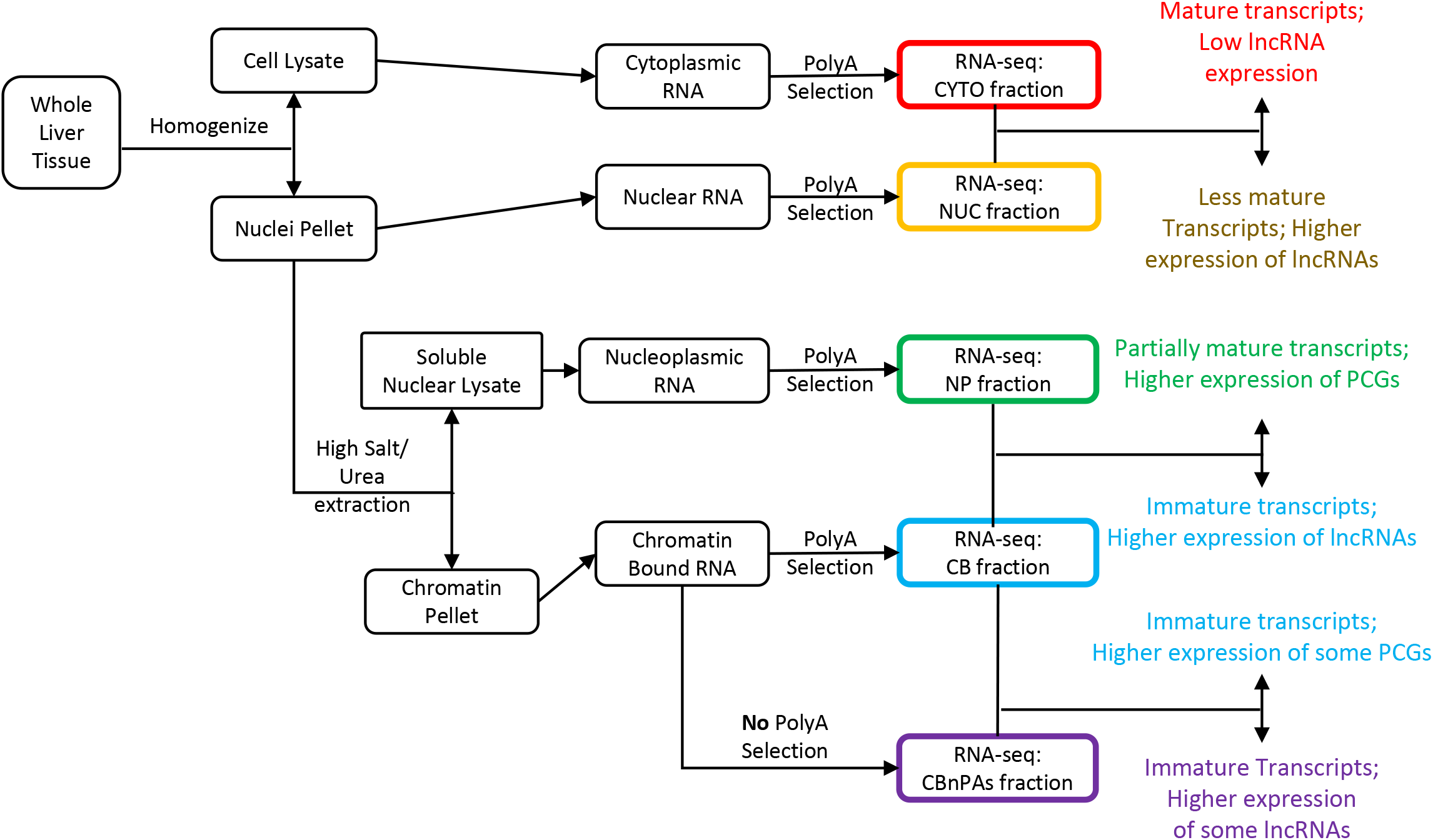
Schematic outline of the purification of mouse liver RNA from liver subcellular fractions. From left to right, liver tissue frozen at -80C was homogenized as described in Methods then centrifuged to separate nuclei from the cellular lysate, which were respectively used to purify cytoplasmic and nuclear RNA. Nuclei were extracted with urea and high salt to obtain a nuclear lysate and chromatin pellet, which were used to purify the corresponding two RNA fractions. RNA isolated from each of the four subcellular fractions (cytoplasm, nucleus, nucleoplasm, chromatin-bound) was polyA-selected and Illumina RNA-seq libraries then prepared. Additionally, a non-polyA-selected RNA-seq Library was prepared from the chromatin-bound fraction. Differential expression analysis was performed using edgeR for the three pairwise comparisons shown at the right: cytoplasm vs nucleus, nucleoplasm vs chromatin-bound, and chromatin-bound (polyA-selected) vs chromatin-bound (non-polyA-selected) to identify lncRNA and PCG transcripts significantly enriched in each subcellular fraction.

**Fig. S2.**
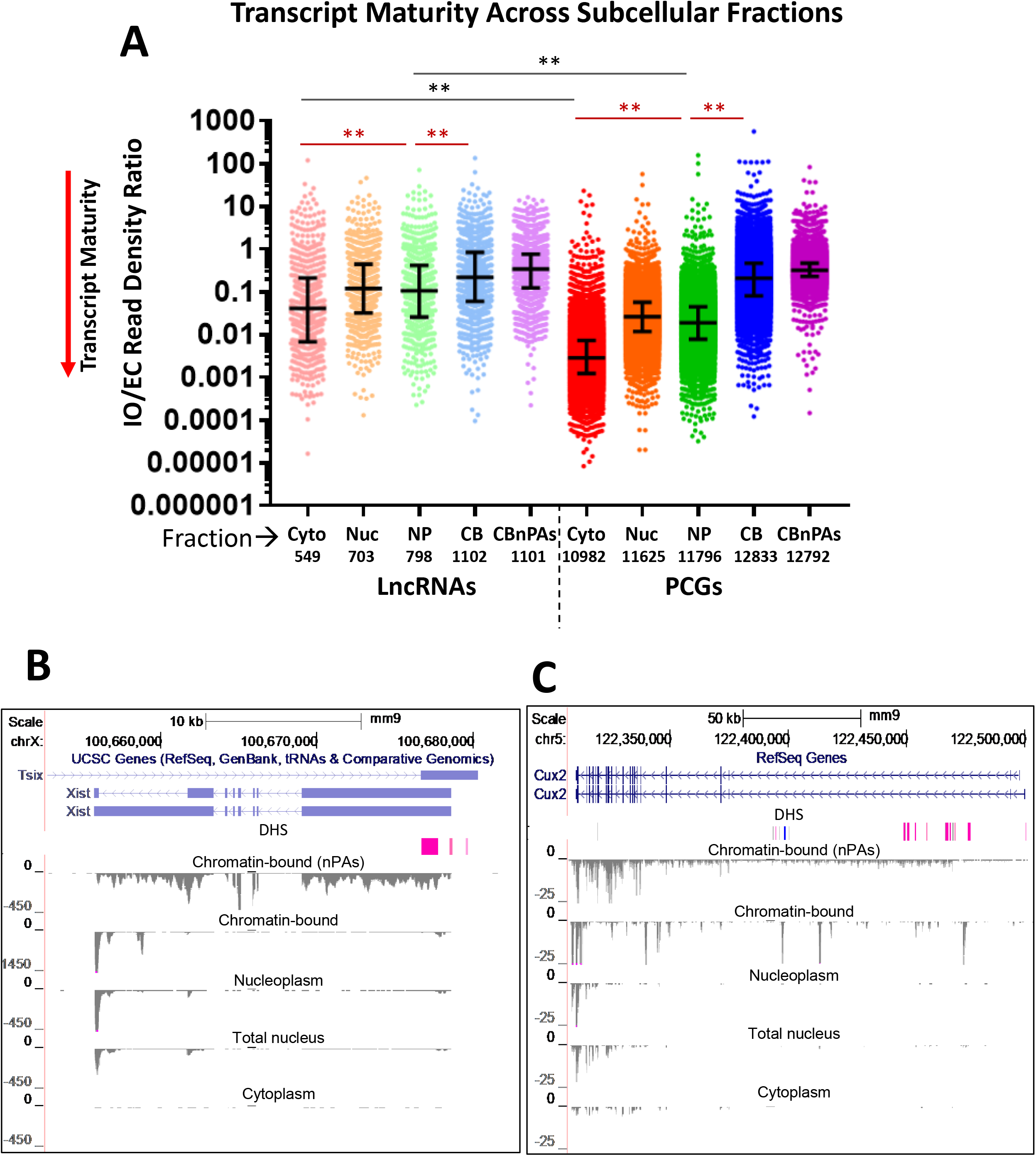
Transcript maturity across subcellular fractions by analysis of IO/EC read density ratios. The distribution of IO/EC read density ratios for individual genes, calculated by dividing the weighted normalized read density of the intronic only (IO) reads vs the exonic collapsed (EC) reads for 1,442 multi-exonic lncRNAs and 13,737 multi-exonic PCGs expressed in vehicle treated female liver in any fraction (mean of n=3 livers). The number of genes expressed per subcellular fraction is shown under each column: Cytoplasm (Cyto), Nucleus (Nuc), Nucleoplasm (NP), Chromatin-bound (CB), Chromatin-bound non-PolyA selected (CBnPAs). Error bars represent the interquartile range of the distribution from the median (horizontal midline). Black brackets compare lncRNA to PCG ratios within the same fraction and red brackets compare lncRNA ratios, or PCG ratios, between fractions (** = adjusted p-value < 0.0001). See Fig. 2 for corresponding results in vehicle-treated male liver. The underlying data used to generate these graphs are found in Table S1F. (**B**) and (**C**) UCSC Browser screen shot showing BigWig files of minus strand sequence reads for each of the five indicated subcellular fractions for Xist (lnc15394) and Cux2 in untreated female mouse liver. Extensive reads seen across the gene body in the chromatin bound fraction are substantially depleted after polyA-selection (top vs second reads track); however, multiple distinct peaks within intronic regions remain, most notably for Cux2. BigWig Y-axis scales are marked on the left. Both genes show female-specific expression, with many fewer sequence reads in corresponding fractions from male liver (not shown). DHS, DNase hypersensitivity sites, indicating open chromatin. These same patterns were seen in all three biological replicates. DHS showing significantly greater accessibility in female liver are marked in pink, and one male-biased DHS within an intron of Cux2 is marked in blue [100]. Also see Fig. 2.

**Fig. S3.**
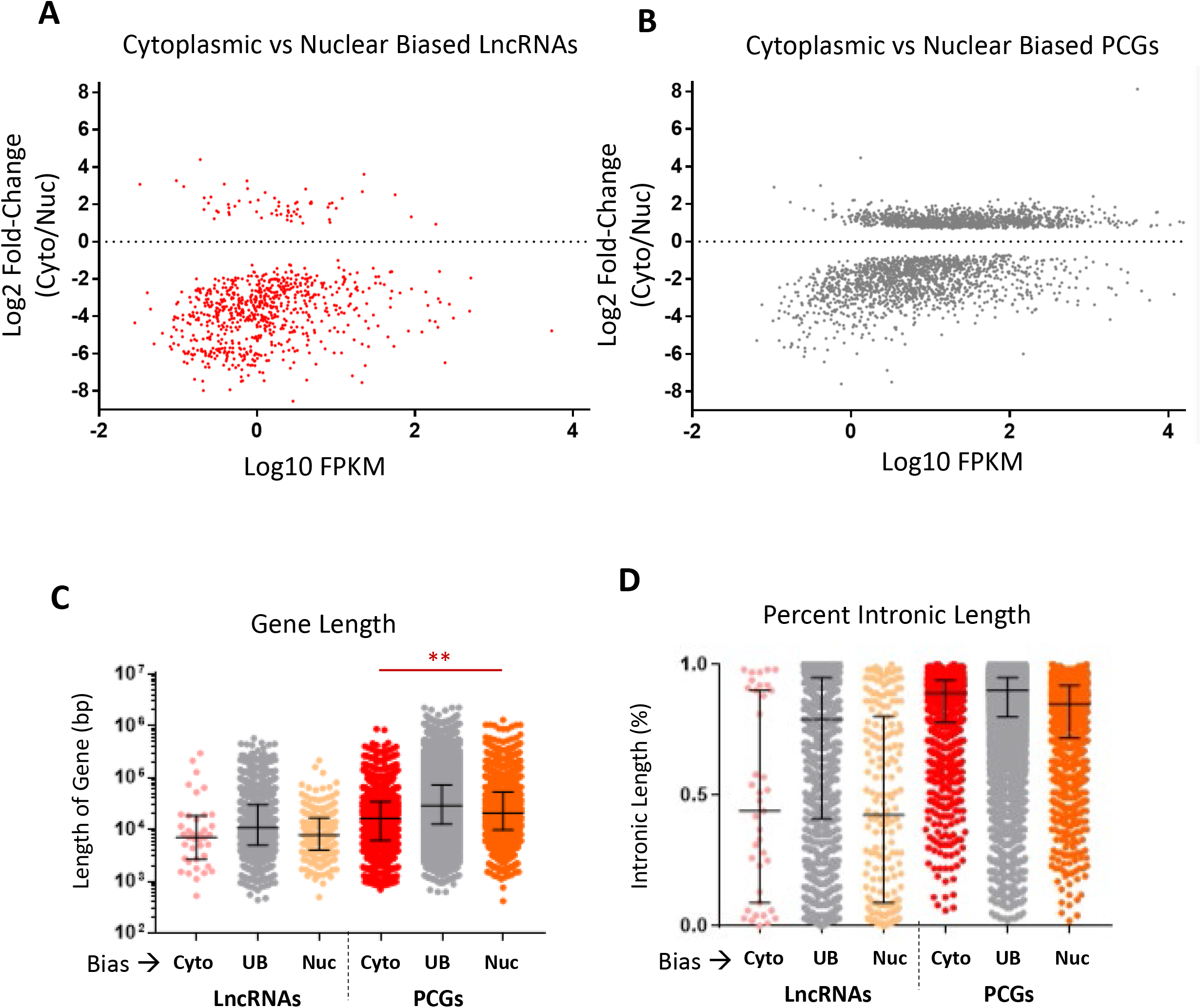
Expression, fold-change and gene length/percent intronic length data for cytoplasmic versus nuclear fractions. Log2 fold-change versus log10 FPKM of cytoplasmic (positive y-axis values) versus nuclear lncRNAs (negative y-axis values) (A) and PCGs (B). Data shown is the same as in Fig. 3A, but presented here as separate graphs for lncRNAs and PCGs to facilitate their visual comparison (also see Table S2A, columns D and E). Comparison of the cytoplasmic biased (Cyto), unbiased (UB) and nuclear biased (Nuc) gene lengths (C) and % intronic length (D) (Table S1D, columns M-Q). A significant difference between the Cyto-biased and Nuc-biased genes was seen for PCGs, but not for lncRNAs, but only in the gene length comparisons. Error bars represent the interquartile range of the distribution from the median (horizontal midline). Red brackets compare lncRNA ratios, or PCG ratios, between sex-biased groups (** = adjusted p-value < 0.0001).

**Fig. S4.**
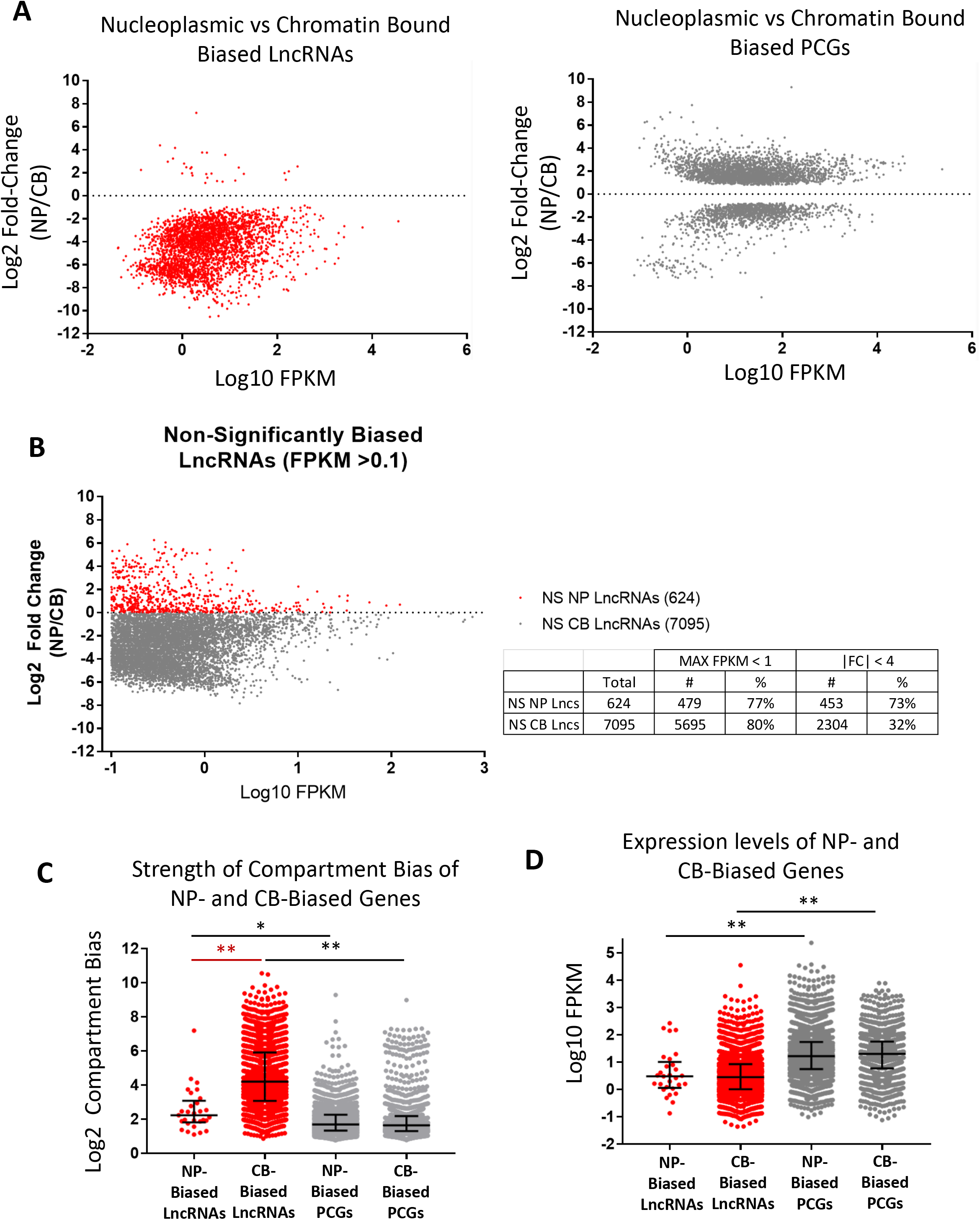
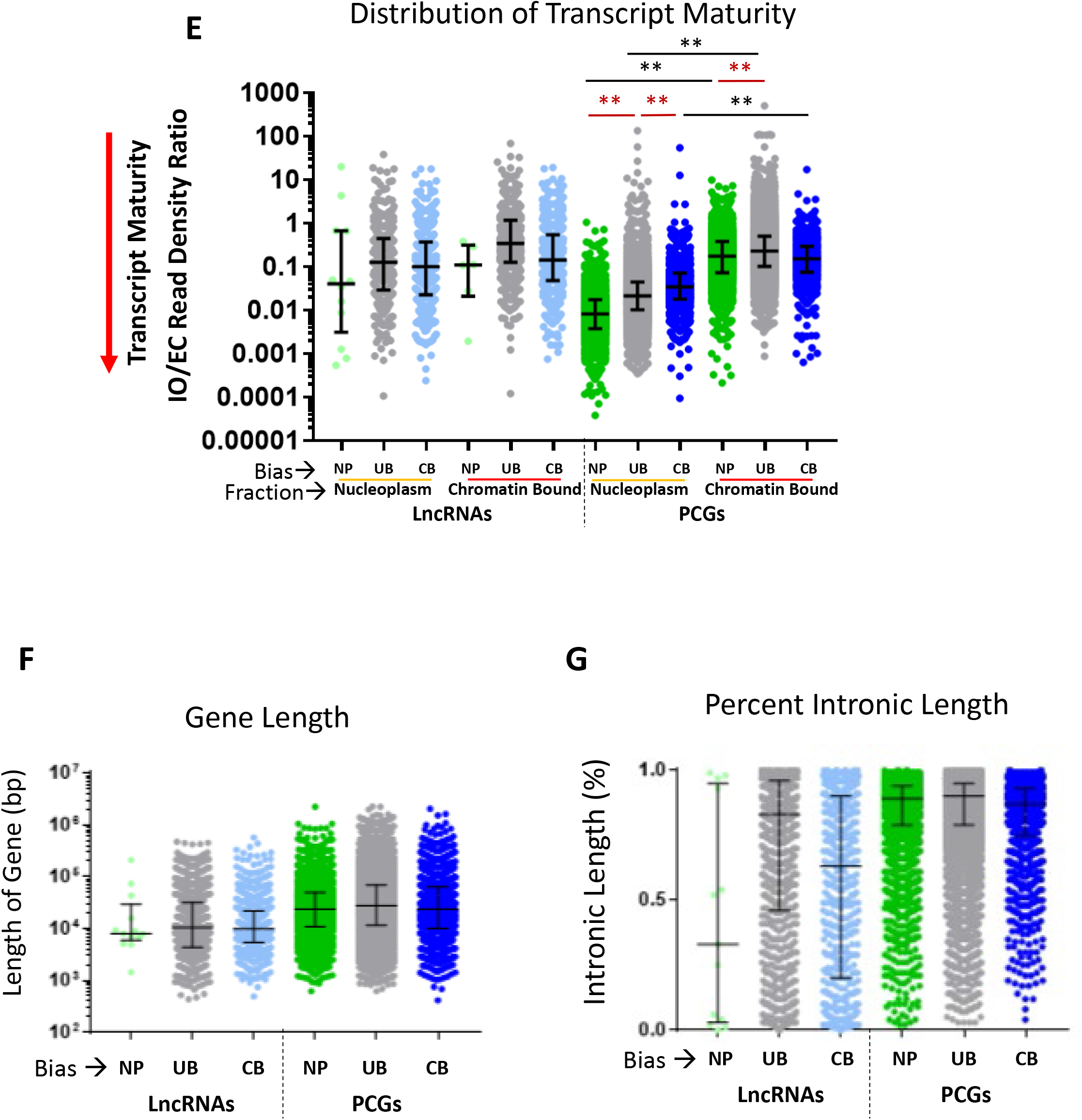
Expression, fold-change, transcript maturity and gene length/percent intronic length for nucleoplasmic versus chromatin-bound fractions. (**A**) Log2 fold-change versus log10 FPKM of nucleoplasmic (positive y-axis values) versus chromatin-bound (negative y-axis values) for lncRNAs (left) and PCGs (right). Data shown is the same as in Fig. S5A, but presented here as separate graphs for lncRNAs and PCGs to facilitate their visual comparison (also see Table S2B, columns D and E). (**B**) Shown are log2 fold-change versus log10 FPKM values, as in A, for 7,719 other lncRNAs, all expressed at FPKM > 0.1, and which did not meet our stringent criteria of adjusted p-value < 0.001 for chromatin or nucleoplasmic bias. 7,095 (92%) of these lncRNAs were more abundant in the chromatin-bound fraction (Table S2B). In many cases, the bias of these lncRNAs for the chromatin-bound fraction was very strong (y-axis values down to log2 fold-change < -4) but missed our stringent significance threshold owing to their low expression (∼80% at FPKM < 1). (**C**) Strength of compartment bias and (**D**) distributions of FPKM expression levels for lncRNAs and PCGs, grouped based on their nucleoplasmic (NP) or chromatin-bound fraction bias (CB). (**E**) Transcript maturity (IO/EC read density ratio, Table S1F) is graphed for the nucleoplasmic and chromatin-bound compartments for the lncRNAs and PCGs that show significant nucleoplasmic bias (NP) or chromatin-bound fraction bias (CB), or are unbiased (UB) with regards to these two compartments. Only multi-exonic genes were included in this analysis. There was no significant difference between NP, UB or CB lncRNAs between the NP and CB compartments. (F and G), comparison of the NP, UB, and CB gene lengths (**F**) and % intronic length (**G**) (see Table S1D, columns M-Q). There were no significant differences between three types of biased genes for either lncRNAs or PCGs with respect to gene length and % intronic length. Error bars represent the interquartile range of the distribution from the median (horizontal midline). Black brackets compare lncRNAs to PCGs within the same fraction, and red brackets compare lncRNAs, or PCGs, between fractions. *= adjusted p-value < 0.05; ** = adjusted p-value < 0.0001.

**Fig. S5.**
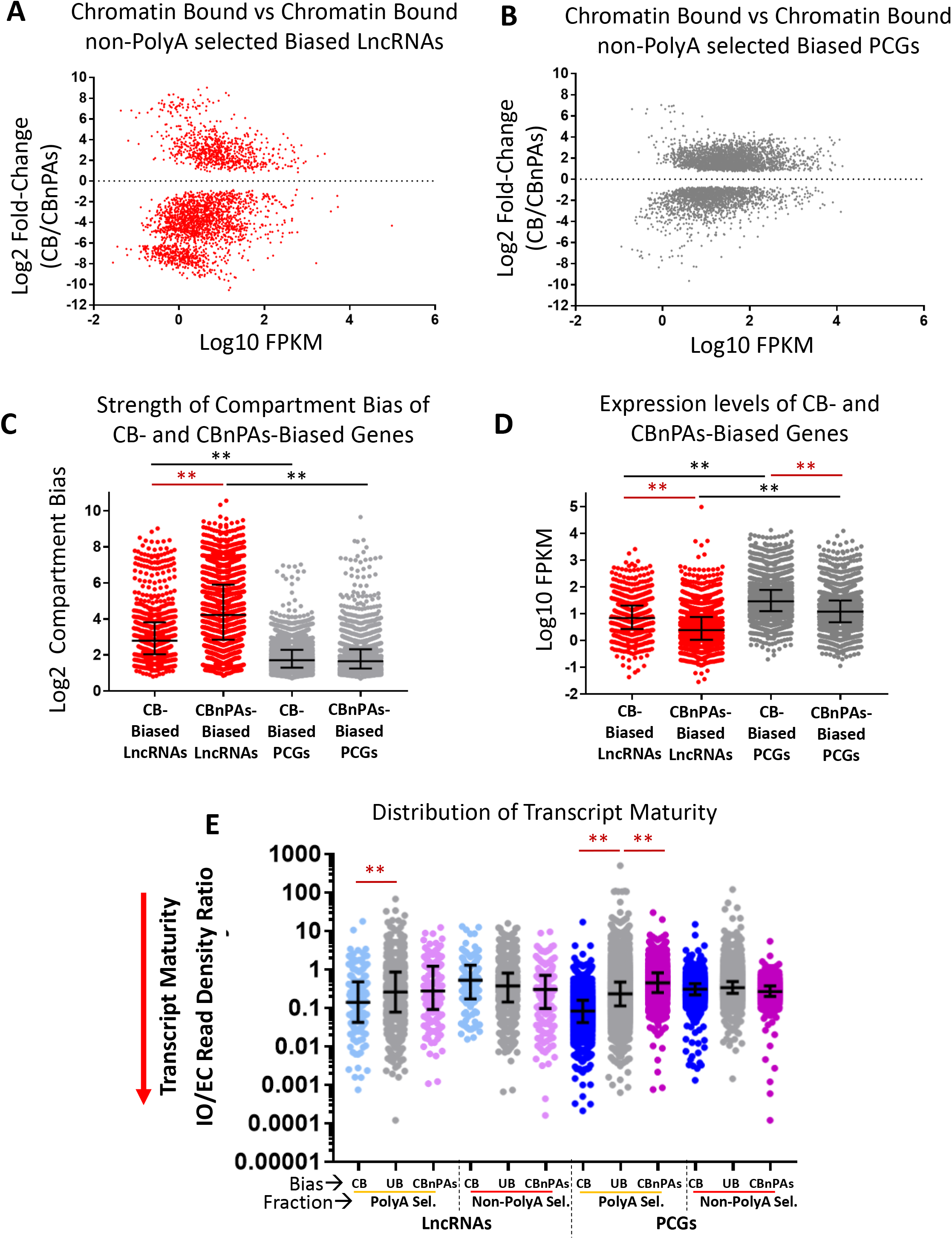
Expression, fold-change and transcript maturity in the chromatin bound versus chromatin bound non-polyA selected fractions. Log2 fold-change versus log10 FPKM of chromatin-bound (positive y-axis values) versus chromatin-bound non-polyA selected lncRNAs (negative y-axis values) (**A**) and PCGs (**B**), presented here as separate graphs (c.f., combined graph in Fig. 4B. Also see Table S2C, columns D and E. Strength of compartment bias data (**C**) and FPKM expression level (**D**) for lncRNAs and PCGs that show chromatin-bound (CB) and chromatin-bound non-polyA selected (CBnPAs) bias. (**E**) Transcript maturity data (IO/EC read density ratio, Table S1F) is graphed in the chromatin-bound and chromatin-bound non-polyA selected fractions for those lncRNAs and PCGs that are biased for the chromatin-bound (CB) or chromatin-bound non-polyA biased (CBnPAs) unbiased fraction (UB) or are unbiased (UB). Only multi-exonic genes were included in this analysis, and only differences between the CB and UB and UB and CBnPAs groups were evaluated for statistical significance in the PolyA selected fraction. Error bars represent the interquartile range of the distribution from the median (horizontal midline). Black brackets compare lncRNAs to PCGs within the same fraction, and red brackets compare lncRNAs, or PCGs, between fractions (*= adjusted p-value < 0.05; ** = adjusted p-value < 0.0001).

**Fig. S6.**
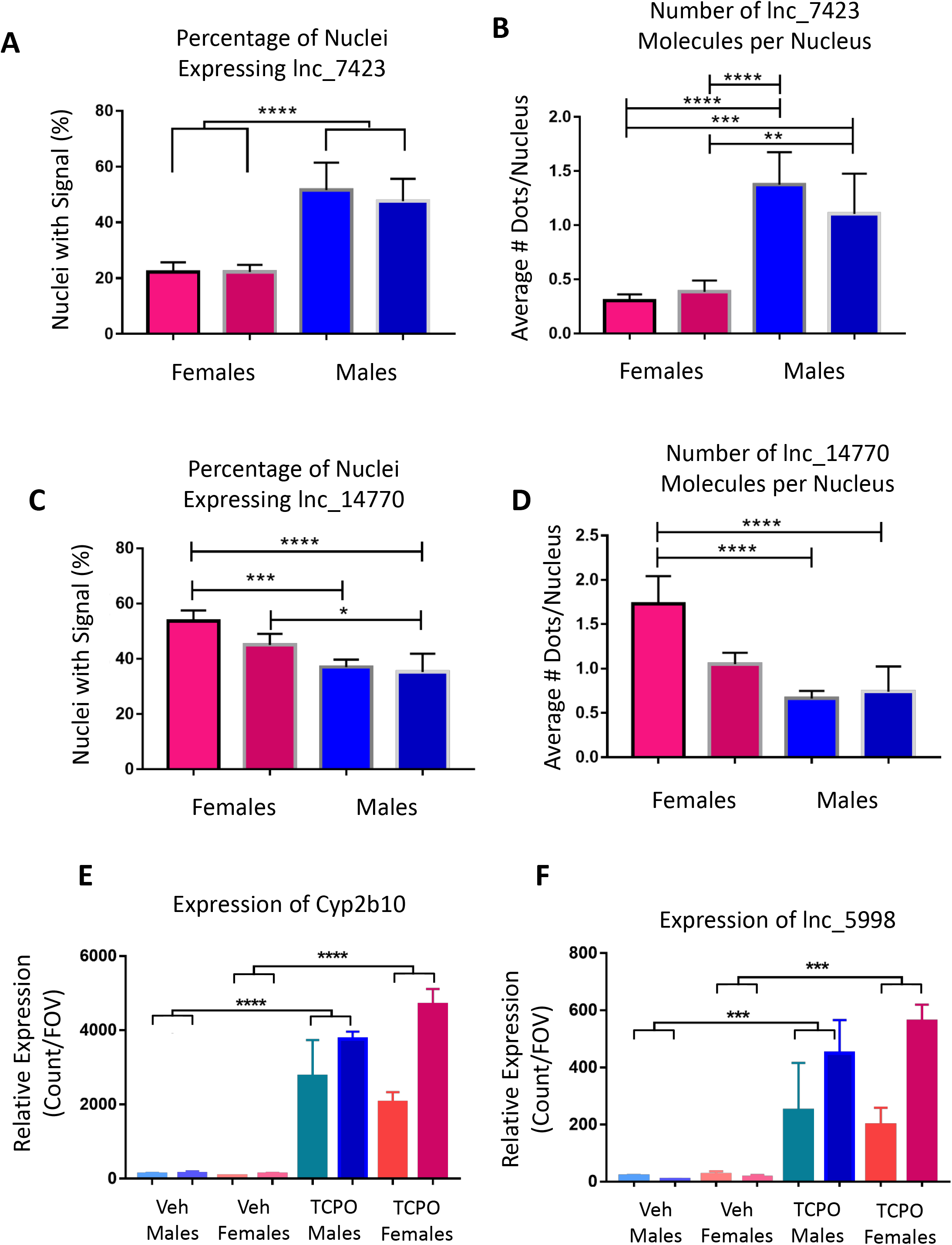
Quantification of lncRNA and PCG expression from smFiSH analysis. Quantification of the numbers of nuclei expressing lnc_7423 (A) and the number of lnc_7423 molecules per nucleus (B) in two individual female livers and two individual male livers. Mean percentage or counts are graphed based on five fields of view for each liver, and error bars represent the SEM. Quantification of the number of nuclei expressing lnc_14770 (C) and the number of lnc_14770 molecules per nucleus (D) in two female livers and two male livers. Mean percentage or counts are graphed for five fields of view for each liver, and error bars represent the SEM. Relative expression of Cyp2b10 (E) and lnc_5998 (F) in male or female liver, either untreated (UT) or from mice treated with TCPOBOP for 51 hr (TCPO). Mean counts are graphed for 5 fields of view for each liver, and error bars represent the SEM. Asterisks show the significance of student t-tests (p-values: * < 0.05, ** < 0.01, *** < 0.001, **** < 0.0001).

